# Context-Invariant Encoding of Reward Location in a Distinct Hippocampal Population

**DOI:** 10.1101/207043

**Authors:** Jeffrey L. Gauthier, David W. Tank

**Affiliations:** Princeton Neuroscience Institute, Princeton University, Princeton, NJ

## Abstract

The hippocampus plays a critical role in goal-directed navigation. Across different environments, however, the hippocampal map seems to be randomized, making it unclear how goal locations might be encoded. To address this question, we trained mice to seek reward at several locations on virtual linear tracks, and optical recordings of calcium activity were made at cellular resolution from two major hippocampal output structures, CA1 and the subiculum. These experiments revealed a population of neurons that were consistently active near reward locations. Their pattern of activity sharply contrasted with simultaneously-recorded place cells and even persisted across environments, when other cells remapped randomly. Their timing was closely correlated with reward anticipation behaviors, yet could not be explained by the behaviors themselves, raising the possibility that reward-associated cell signals were used by mice to identify the reward location. These results demonstrate that the hippocampus employs a context-invariant marker of goal locations, and reveal a novel target for studying how the hippocampus contributes to navigation.

## 1. Introduction

The hippocampus is crucial for many kinds of spatial memory [12, 22, 7], and in particular learning to navigate to an unmarked goal location [29, 15, 40]. Consistent with this role, individual hippocampal neurons exhibit spatially-modulated activity fields, or place fields, that encode the animal’s current location [33], and collectively form a map-like representation of space [35]. In each environment an animal explores, the hippocampal map is sensitive to the particular constellation of features experienced in that environment: the geometry of boundaries [31], visual features [24], odors [3], and even the absolute time when it was explored [41]. While context-specific representations likely benefit other hippocampal functions, such as episodic memory [7], they seem poorly suited to guide goal-directed navigation. In each new environment, the hippocampal encoding is essentially randomized [24], meaning any downstream circuit sampling from the population must learn a new, idiosyncratic code to localize the goal.

There is some evidence, however, that goals might be represented differently than other points in space. In the CA1 region of the hippocampus, place fields often occur at higher density near goal locations than other parts of the environment [15, 19, 13, 9, 43]. We therefore sought to identify whether any cells contributing to the excess density might be specialized to encode the goal location, similar to other hippocampal neurons that seem to encode abstract categories [38, 25].

This question was focused on CA1 and subiculum, two major outputs of the hippocampal formation [2]. Because cells composing the excess density are a relatively small proportion of place cells (~ 15%), and because previous studies have reported low yield from electrode recordings in the subiculum [47, 21], optical imaging was used to record activity in transgenic mice expressing the calcium indicator GCaMP3 [39]. Mice were trained to seek rewards at different locations within several virtual reality environments, and the activity of thousands of individual neurons was tracked to identify whether any were specialized to encode reward.

## 2. Results

### 2.1. Moving Reward Location Within One Environment

Water-restricted mice were trained to traverse a virtual reality environment in an enclosure that allowed simultaneous two-photon imaging at cellular resolution [17, 13, 14]. The virtual environment was a linear track with a variety of wall textures and colors that provided a unique visual scene at each point (Figure 1b). When mice reached a fixed location on the track (366 cm), a small water reward was delivered from a tube that was always present near the mouth. After the end of the track, the same pattern of visual features and reward delivery was repeated, creating the impression of an infinite repeating corridor.

**Figure 1.**
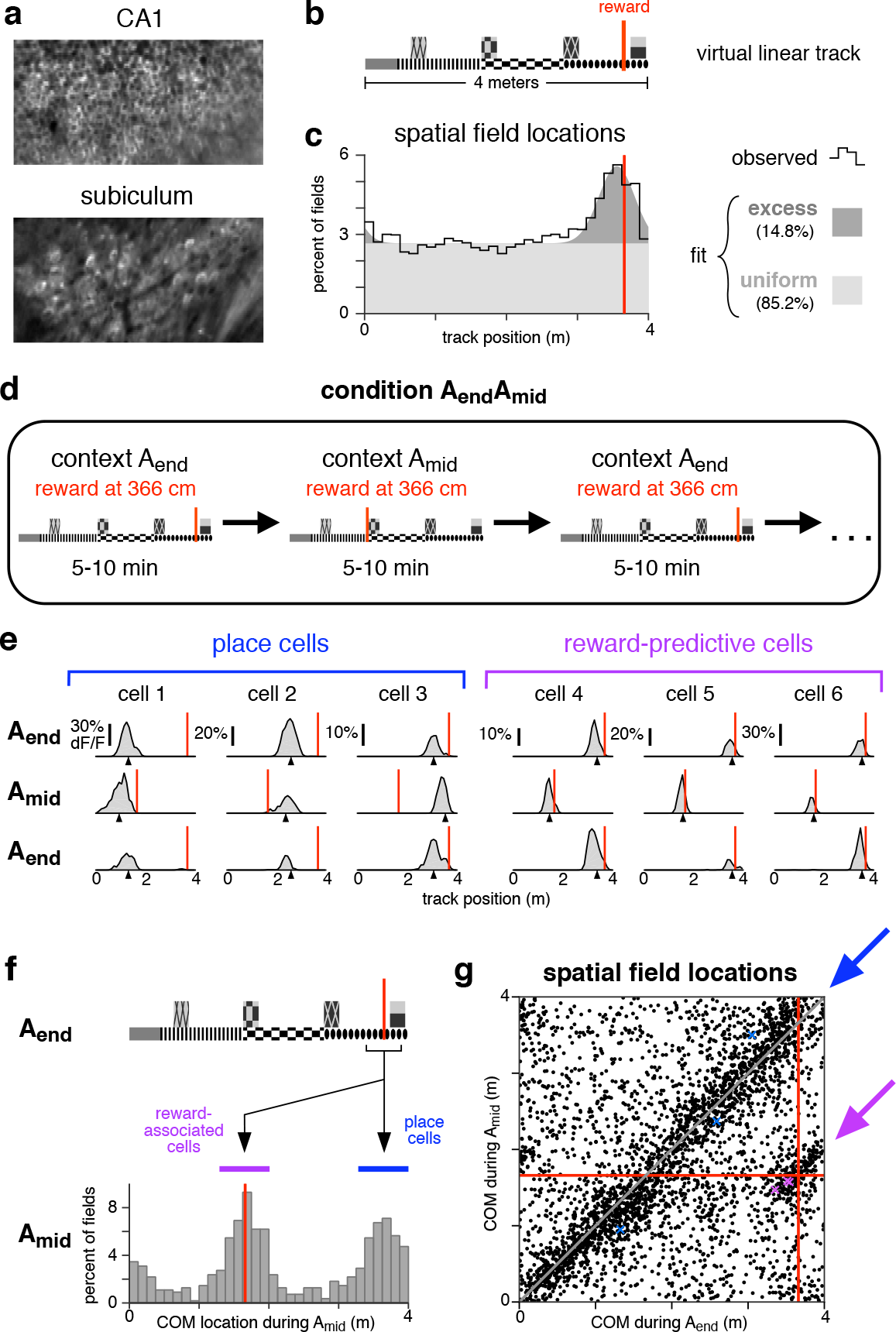
A distinct population of hippocampal neurons are consistently active near reward. **a.** Typical fields of view in CA1 and subiculum of neurons expressing GCaMP3. Image widths are 200 um. **b.** Schematic of the virtual linear track and reward location (red line). **c.** COM locations of all cells with a spatial field during condition A_end_ (9,761 cells, 11 mice). Black line shows observed density, gray patches show density of a fitted mixture distribution consisting of a uniform distribution (light gray) and a Gaussian distribution (dark gray, mean 355 cm, s.d. 25 cm). **d.** Schematic of condition in which reward location shifted between two locations. **e.** Activity of six simultaneously-recorded CA1 neurons during the first three blocks of one session of condition A_end_A_mid_. Each column shows the spatially-averaged activity of one cell in the first (top), second (middle), and third (bottom) blocks. Activity on each traversal was spatially binned (width 10 cm), filtered (Gaussian kernel, radius 10 cm) and averaged (70th percentile) across all traversals, excepting the first three traversals of each block. Black arrowheads indicate COM location computed by pooling trials from all blocks of a single context (A_end_ or A_mid_, see Methods). Red lines indicate reward location in each block. **f.** Top: track diagram. Bottom: COM locations during A_mid_ of cells with a spatially-modulated field located within 25 cm of reward during A_end_ (square bracket beneath track diagram, 1,171 cells). Red lines indicate reward location, colored bands indicate clusters of reward-associated cells (purple) or cells whose field remained in the same location (blue). **g.** The COM locations of all cells with spatial fields during both A_end_ and A_mid_ (3,761 cells). Red lines indicate reward location. Arrows indicate regions defining reward-associated cells (purple) and place cells with stable field locations (blue).

While mice interacted with the virtual environment, optical recordings of neural activity were made in CA1 and subiculum (Figure 1a). Consistent with previous studies in real and virtual environments [35, 13, 4], many neurons in both regions exhibited place fields, i.e. activity patterns that were significantly modulated by position on the track (see Table 1). The activity location of each spatially-modulated cell was summarized by the center of mass (COM) of its activity averaged across trials.

**Table 1.**
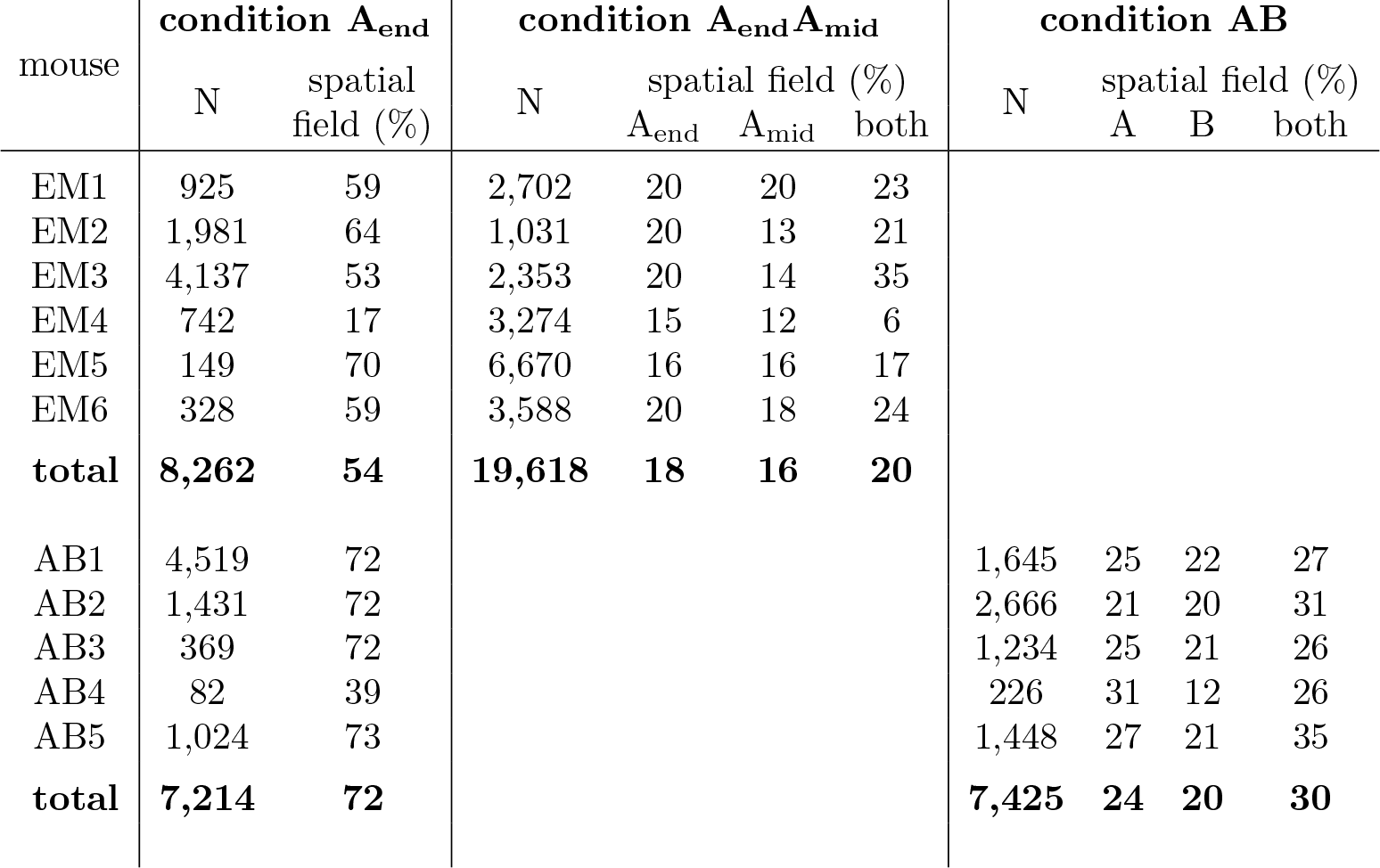
Recorded cell counts for each mouse used in population analyses (EM1-6 for condition A_end_A_mid_, AB1-5 for condition AB), and the fraction of recorded cells with a spatially-modulated field. Note that the recording and cell finding techniques were likely biased towards detecting more active neurons, potentially increasing the fraction of cells with a spatial field compared to estimates from other techniques.

| mouse | condition $A_{\text{end}}$ | | condition $A_{\text{end}}A_{\text{mid}}$ | | | | condition AB | | | |
| --- | --- | --- | --- | --- | --- | --- | --- | --- | --- | --- |
| | N | spatial field (%) | N | spatial field (%)<br>$A_{\text{end}}$ | $A_{\text{mid}}$ | both | N | spatial field (%)<br>A | B | both |
| EM1 | 925 | 59 | 2,702 | 20 | 20 | 23 |  |  |  |  |
| EM2 | 1,981 | 64 | 1,031 | 20 | 13 | 21 |  |  |  |  |
| EM3 | 4,137 | 53 | 2,353 | 20 | 14 | 35 |  |  |  |  |
| EM4 | 742 | 17 | 3,274 | 15 | 12 | 6 |  |  |  |  |
| EM5 | 149 | 70 | 6,670 | 16 | 16 | 17 |  |  |  |  |
| EM6 | 328 | 59 | 3,588 | 20 | 18 | 24 |  |  |  |  |
| **total** | **8,262** | **54** | **19,618** | **18** | **16** | **20** |  |  |  |  |
| AB1 | 4,519 | 72 |  |  |  |  | 1,645 | 25 | 22 | 27 |
| AB2 | 1,431 | 72 |  |  |  |  | 2,666 | 21 | 20 | 31 |
| AB3 | 369 | 72 |  |  |  |  | 1,234 | 25 | 21 | 26 |
| AB4 | 82 | 39 |  |  |  |  | 226 | 31 | 12 | 26 |
| AB5 | 1,024 | 73 |  |  |  |  | 1,448 | 27 | 21 | 35 |
| **total** | **7,214** | **72** |  |  |  |  | **7,425** | **24** | **20** | **30** |

The COMs of spatial fields were distributed throughout most of the track at uniform density [32, 34], but an excess density was located near the reward, both when pooling all cells (Figure 1c) and considering CA1 and subiculum separately (Supplemental Figure 1). This enhancement of the representation near reward was consistent with previous studies in both real and virtual environments [15, 19, 13, 9], and it permitted subsequent experiments to characterize the cells composing the increased density.

Two possibilities were considered. First, the excess fields might have reflected an increased number of place fields, ,i.e. fields encoding a particular position on the track as defined by visual landmarks, They might have formed at a higher rate near reward due to the salience of the location, increased occupancy time, or other factors [18]. Alternatively, some excess fields might have arisen from cells that did not encode a place per se, but instead were specialized to be active near reward [6]. To distinguish these possibilities, the reward was moved to a different part of the environment (condition A_end_A_mid_, Figure 1d).

During reward-location alternation sessions, many cells exhibited spatial fields (Table 1), and the fields of most cells remained in the same location. At the same time, the fields of some cells shifted to match the reward location. These two behaviors are illustrated for one session (Figure 1e): stable spatial fields were observed throughout the track (cells 1-3), while a separate population shifted to be consistently located near reward (cells 4-6).

Stability versus shifting to track the reward was a discrete difference, which can be appreciated by considering how cells near one reward location remapped when the reward shifted. Of cells active near the reward during A_end_ (Figure 1f, black bracket), those with spatial fields during A_mid_ tended to be found in two locations: either near the same part of the track (blue band), or immediately before the reward location at 166 cm (purple band). This pattern was significantly bimodal (p < 1e-5, Hartigan’s Dip Test), indicating that cells associated with reward formed a discrete subgroup, distinct from those remaining active near the same visual landmarks.

To identify response types in the entire population, remapping was characterized for all cells with a spatial field in both contexts (Figure 1g). When the reward shifted, the fields of most cells remained in the same location (blue arrow), while a separate population shifted to be consistently active near reward (purple arrow). The latter group will be referred to as “reward-associated cells”, and they composed 4.2% of cells with fields in both conditions, or 0.8% of all recorded cells. Of the remaining cells, five response types were observed: cells with no spatial field in either condition (46.8% of all recorded cells), cells with a field in A_end_ only (17.8%), cells with a field in A_mid_ only (15.8%), cells with a field in both contexts that shifted by less than 50 cm (10.9%), and cells with a field in both contexts that remapped to new, apparently random locations (7.9%). These response types are consistent with the well-characterized physiology previously described in CA1, the subiculum, and throughout the hippocampal formation [2] and they will be referred to as “place cells”. Against this backdrop, reward-associated cells stood out as a distinct population.

An additional important distinction can be made among reward-associated cells: some were active prior to reward delivery, and others were active at a location subsequent to the reward. While both categories might be relevant for navigation, cells active before reward are particularly noteworthy, since their activity could not be explained as a response to the immediate visual environment, and subsequent analyses will show that they were not responding to reward anticipation behaviors. Instead, the remapping of cells active before reward seemed to incorporate recall of past events, i.e. at which site the reward had recently been delivered. These cells will be referred to as “reward-predictive cells”, in the sense that their field locations reliably indicated where the reward would be delivered, even before it arrived. The name is not intended to suggest they necessarily play a role in memory or reward prediction behaviors, though these possibilities will be considered.

### 2.2. Switching Between Two Environments

If reward-associated cells were truly specialized to encode reward, downstream circuits would likely benefit from those signals arising from same cell population in each new environment. Contrary to the consistency this scheme requires, hippocampal representations of different contexts are highly divergent [24, 27]. We therefore asked whether reward-associated activity was randomly re-assigned to different cells in a new environment, or if instead it was produced by the same cells.

To distinguish these possibilities, a separate cohort of mice was trained on a new paradigm, condition AB, in which mice alternated block-wise between the original track (track A), and a second, shorter track with distinct visual textures (track B). On track B, reward was also delivered near the end, and on both tracks spatial fields occurred at increased density near reward (Supplemental Figure 2a).

As mice alternated between tracks A and B, the fields of some cells shifted to different, apparently random locations, while others were consistently active near reward, indicating that reward-associated cells formed a separate group. These response types are illustrated for several cells in Figure 2b, and they were shown to be representative in several population-level analyses.

**Figure 2.**
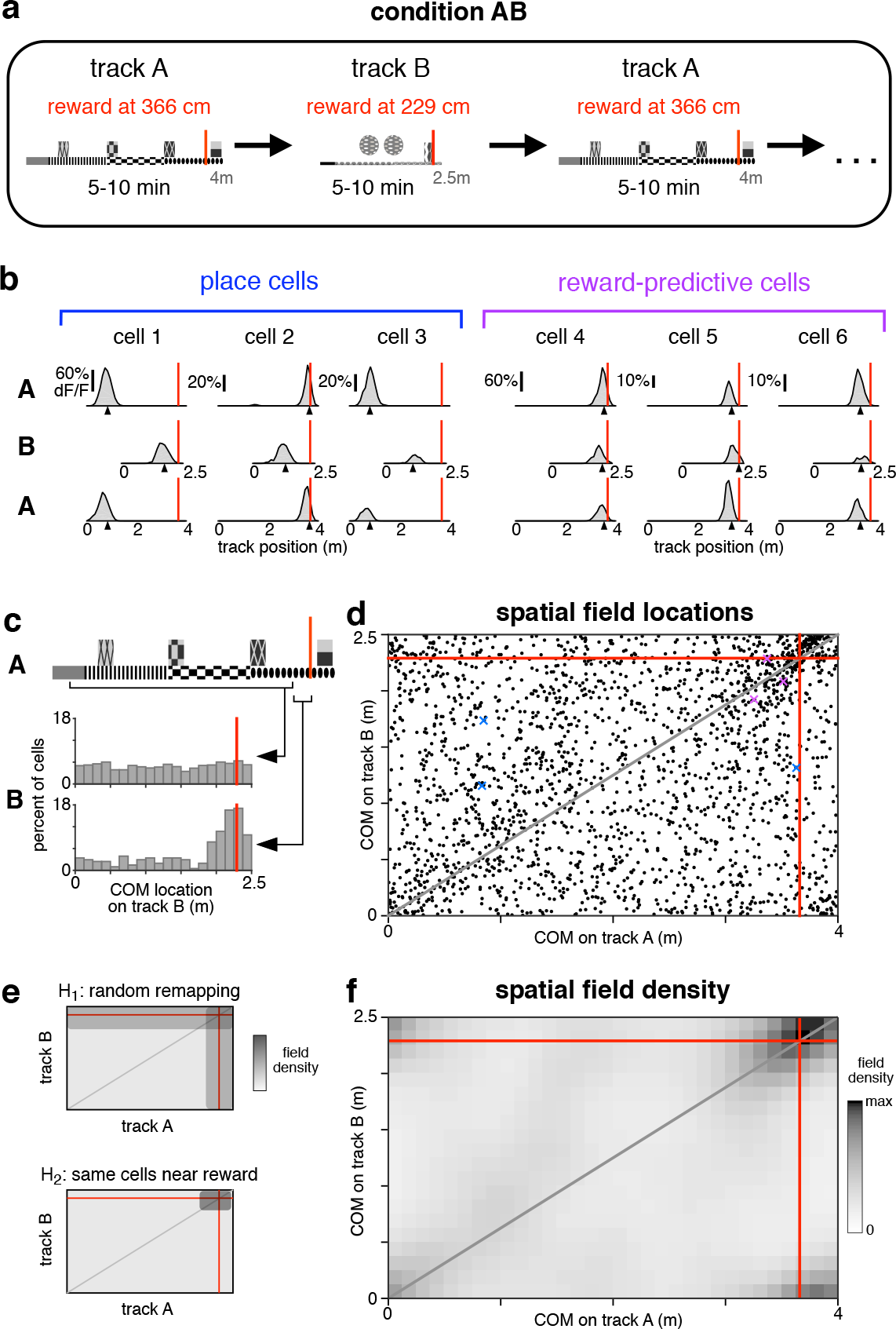
Reward-associated cell identity persists across contexts. **a.** Schematic of condition in which mice were teleported between two different virtual linear tracks. **b.** Activity of six simultaneously-recorded CA1 neurons during the first three blocks of one session of condition AB. Same conventions and spatial-averaging procedure as in Figure 1e, except that all traversals were included. **c.** COM locations on track B for two populations of cells. Upper histogram: cells with a spatial field on track A located between 25 cm after track start to 25 cm before reward (wide square bracket). Lower histogram: cells with a spatial field on track A located in the 25 cm preceding reward (narrow square bracket). **d.** COM locations of all cells with a spatial field on both track A and track B (2,168 cells, 5 mice), red lines indicate reward locations. Gray line indicates proportionally equivalent locations on the two tracks. **e.** Schematic of COM density under two hypotheses for how spatial fields remapped (see text). **f.** Observed density of COMs on track A and track B (same data as in d), spatially-binned (width 12.5 cm) and smoothed (2D Gaussian kernel, radius 20 cm).

Across the tracks, most spatial fields shifted in a manner consistent with global remapping. Field locations (Figure 2d) spanned the complete range of possible remappings, and their density appeared approximately uniform, possibly excepting regions near reward. This effect was confirmed quantitatively by comparing to a previous study that employed similar recording techniques [9]. That study found the average population vector correlation across two distinct physical treadmills was 0.22, while in the present study the equivalent value was 0.099 (95% confidence interval 0.094 to 0.103). This comparison confirmed that switching between tracks A and B elicited global remapping, and provided further evidence that virtual environments are capable of reproducing much of the same hippocampal phenomenology that has been characterized using other methods [17, 13, 14, 4].

Despite random remapping among most neurons, reward-associated fields appeared to be produced by largely the same cells on both tracks. This was first apparent by comparing remapping in two subsets of cells (Figure 2c). While most place cells remapped to random locations (wide bracket, upper histogram), reward-associated cells on track A tended to be active near reward on track B (narrow bracket, lower histogram).

To demonstrate this effect statistically, quantitative hypotheses were generated for how switching between tracks impacted field locations. The first hypothesis, H1, postulated that cell identities were randomly shuffled between the two conditions, though on each track there was still an increased field density near reward. Under H1, a given cell could, for example, contribute to the excess density of fields near reward in one environment, and then encode a random place on the track in the other environment. The second hypothesis, H2, postulated that cell identities were perfectly preserved. Under H2, reward-associated cells would continue to maintain fields near the reward on both tracks. Meanwhile, place cells would remap randomly, including to locations near the reward, but with the same probability as at other points on the track. Each of these hypotheses predicted a particular distribution for the density of field locations on the two tracks (Figure 2e). To account for a partial contribution of each model, a weighted combination of the H1 and H2 distributions was fit, and it also included cells with a spatial field on only one track (see Methods).

The observed density of field locations (Figure 2f) exhibited an approximately uniform density everywhere, except for an increased density at the intersection of reward locations. Notably, there is no increase in density along the reward lines as predicted by H1, suggesting the data were entirely accounted for by H2. This qualitative impression was confirmed by a numerical fit. Among cells composing the excess density, all remapped according to H2 (100.0%, 95% confidence interval 99.6%-100.0%). The same result was found when CA1 and subiculum were analyzed separately (Supplemental Figure 2). Reward-associated cells with a field on both tracks composed 4.4% of all recorded cells in CA1 and 5.7% in subiculum.

This analysis demonstrated that cell identity was perfectly (100.0%) preserved across the two environments. While the population of place cells remapped to random locations, consistent with the global remapping observed in previous studies, reward-associated cells did not deviate from the reward location. Moreover, reward-associated cells fully accounted for the excess density of fields near reward. This sharp division of cell identities revealed an unexpected degree of consistency in hippocampal encoding of goal locations, and was particularly surprising in CA1, where cell classes that persist across contexts have not been described previously.

### 2.3. Correlation of Reward-Predictive Cells with Reward Anticipation

The previous results showed that reward-predictive cells reliably indicated the reward location, but it was unclear whether mice themselves could predict reward (e.g. anticipatory licking). If so, it would be important to identify whether reward-predictive cells were linked to the behavioral prediction, or if instead they encoded reward location independently of behavior.

In mice that had experienced many traversals of condition A_end_, two kinds of anticipation behaviors were apparent: slowing down prior to reward delivery, and licking the reward tube (Figure 3a). Slowing was typically initiated prior to licking (Supplemental Figure 3), making it the earliest reliable indicator of reward anticipation. In addition, slowing was observed more frequently than licking (not shown), and therefore slowing was used to indicate the onset of reward anticipation.

**Figure 3.**
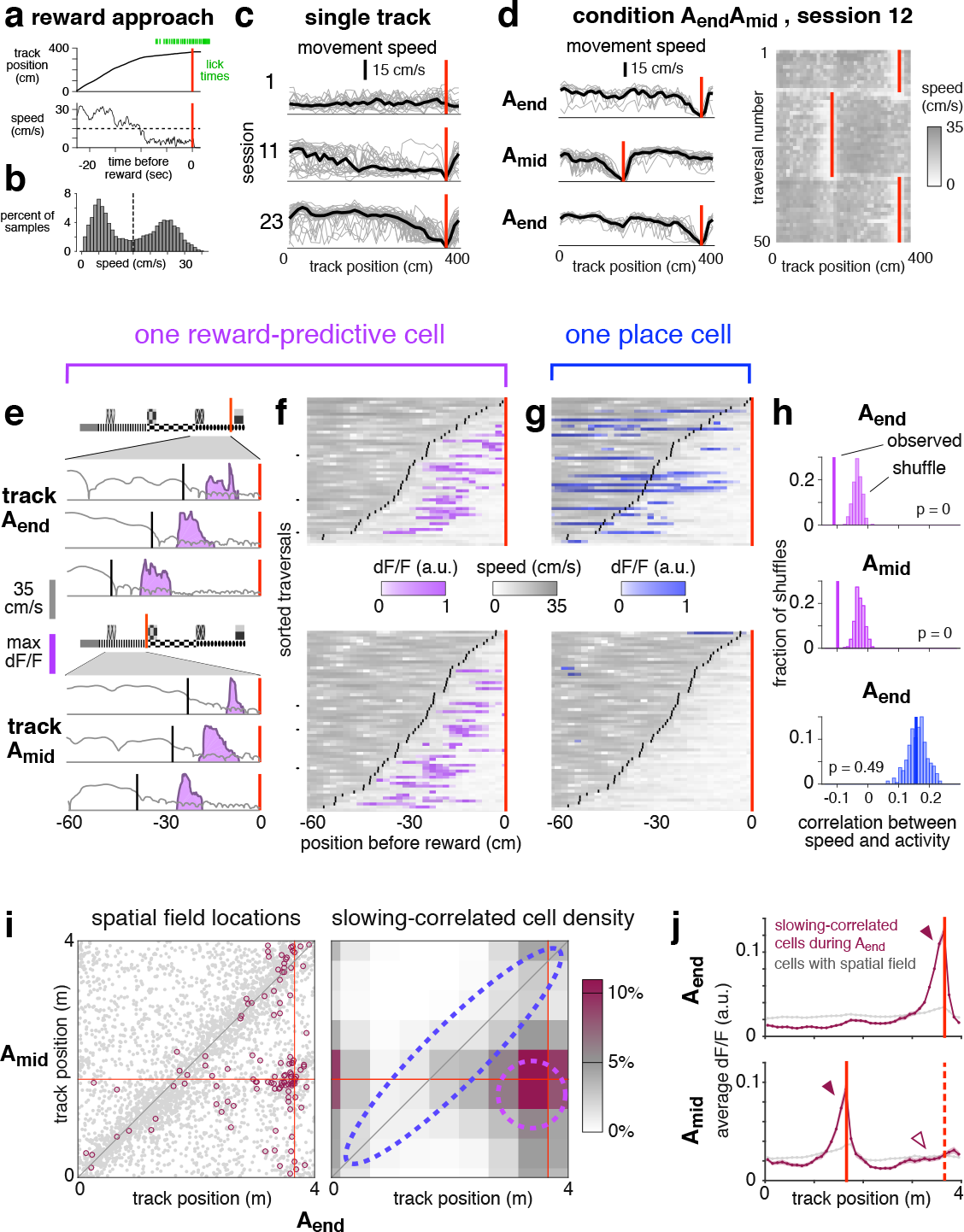
Reward-predictive cell activity is correlated with anticipation of reward. **a.** Representative example of reward approach behaviors. **b.** Movement speeds during one session with slowing threshold (dashed line). **c.** Spatially-binned speed for single trials (gray lines) and averaged across trials (black lines). Red lines indicate reward. **d.** Left: spatially-binned speed for the first three blocks of one session of condition A_end_A_mid_, first block is top panel, same conventions as in b. The first three traversals of each block are omitted. Right: Running speed on the first fifty traversals. **e.** Reward approach behavior on six trials from the session depicted in **c** comparing speed (gray), slowing onset (black), and activity of one reward-predictive cell in CA1 (purple). **f.** Speed (gray) and activity (purple) on all traversals in which slowing onset (black lines) occurred within 6Q cm before the reward location (red line) for same cell as in e. Each pixel shows average in a 2 cm spatial bin. Black tick marks show example trials plotted in e. **g.** Activity of a simultaneously-recorded place cell, same conventions as in **e**, except activity is shown in blue. **h.** Statistical test to evaluate correlation between activity and speed for the cells depicted in **f** and **g**. **i.** Left: COM locations of cells with spatial fields in both A_end_ and A_mid_ (gray points, same data as Figure 1g). Highlighted cells (maroon circles, 116 cells) were slowing-correlated during A_end_ (see definition in text). Right: Lower bound of estimated density of slowing-correlated cells, binned by COM location and spatially smoothed (see Methods). Dashed lines indicate approximate boundaries of reward-predictive cells (purple) and stable place cells (blue). **j.** Upper: average activity of 198 slowing-correlated cells (6 mice, maroon trace) during A_end_ blocks, and all spatially-modulated cells recorded simultaneously (7,343 cells, gray trace). Red line indicates A_end_ reward location. Lower: activity of same cells during the interleaved A_mid_ blocks. Red lines indicate reward location for A_mid_ (solid) and A_end_ (dashed). Bands indicate standard error of the mean. For arrowheads, see text.

Slowing behavior developed gradually throughout training, and with sufficient experience was observed on nearly every trial, as illustrated here for one mouse (Figure 3c). Importantly, in condition A_end_A_mid_, which involved shifting reward delivery, the slowing location rapidly adapted to the current reward location, typically within the first 2-3 traversals (Figure 3d). This demonstrated that the mouse understood the reward alternation paradigm, and further showed that slowly walking was a robust phenomenon that could be used to track reward anticipation at single-trial resolution.

For quantitative analyses, slowing onset required a precise definition. Consistent with previous experiments showing a discrete onset of anticipation behaviors [28], movement speeds here were significantly bimodal (Figure 3b, p < 1e-5, Hartigan’s Dip Test). On each trial, the onset of reward anticipation was defined as speed dropping below a mouse-specific threshold for the last time prior to reward delivery.

The timing of reward anticipation was precisely aligned to the activity of many reward-predictive cells, but this was not generally true of place cells. The difference is illustrated for two simultaneously-recorded cells in Figure 3e-f. On a few representative single traversals (panel e), and across all traversals (panel f), the activity of the reward-predictive cell occurred at approximately the same distance after the onset of slowing. In contrast, a simultaneously-recorded place cell exhibited no such correlation (panel g). These visual impressions were confirmed to be significant using a statistical metric, the percentile correlation, in which the observed value was compared to a shuffle distribution (panel h, see Methods).

These examples were representative of all cells recorded during condition A_end_A_mid_. For each cell, the percentile correlation was compared to how the cell remapped (panel i). Cells that were slowing-correlated (defined as a percentile correlation of 5 or less) were primarily those that maintained fields near the reward (dashed purple outline). Although some cells that exhibited spatial fields in the same location across contexts (dashed blue outline) were also slowing-correlated, they did not occur at a rate exceeding chance.

This result was confirmed in a separate, non-parametric analysis that did not explicitly measure cell density. For cells that were slowing-correlated during A_end_ blocks, fluorescence activity was plotted as a function of position by averaging across cells and traversals (Figure 3j, top panel). As expected, most activity was located just prior to the reward (solid arrowhead), while in the general population it was distributed relatively uniformly (gray trace). During the interleaved A_mid_ blocks (bottom panel), the activity peak of slowing-correlated cells shifted to the current reward location (solid arrowhead), showing that many slowing-correlated cells were also reward-predictive cells. However, at the location where the peak had been observed previously there was no increase above baseline (hollow arrowhead), showing that few if any place cells were slowing-correlated. A similar pattern was observed for cells that were slowing-correlated during A_mid_ (Supplemental Figure 4), and also when considering CA1 and subiculum separately (not shown).

These analyses demonstrated that, during condition A_end_A_mid_, correlation with slowing was not a general feature of the hippocampal ensemble. Instead, it was limited exclusively, or almost exclusively, to reward-predictive cells. This further differentiated the responses of reward-predictive cells from place cells, and raised the possibility that they might contribute to anticipation behaviors.

Interestingly, the correlation with slowing was less prevalent during condition AB. Though some slowing-correlated cells seemed to present, they composed a much smaller fraction of the total population (5.4% of cells with sufficient activity were slowing-correlated on track A during condition AB vs 11.2% during A_end_ blocks of condition A_end_A_mid_). The discrepancy showed that reward-predictive cells were not generally aligned to all instances of reward anticipation, but instead their recruitment depended on particular features of the task. In this case, the important difference might have been increased cognitive demand: during condition A_end_A_mid_, anticipating the reward required accurate recall of recent events, whereas during condition AB anticipation could have relied entirely on the immediate visual cues.

### 2.4. Sequential activation of reward-predictive cells

To identify any reproducible structure in the activity patterns of reward-associated cells, their timing was compared across different contexts. This analysis revealed consistent sequential activation, as illustrated for one session in Figure 4a-b. Comparing blocks of A_end_ to A_mid_, cells were active in a similar order, with a significant correlation in their peak activity times (p < 0.0001). Moreover, their timing across contexts was nearly identical, with the peak fluorescence shifting by a median of only 0.4 seconds. This was a remarkably brief offset in light of several factors: the potential for behavioral variability, the temporal uncertainty of calcium imaging methods, and the fact that the sequence spanned more than 6 seconds.

**Figure 4.**
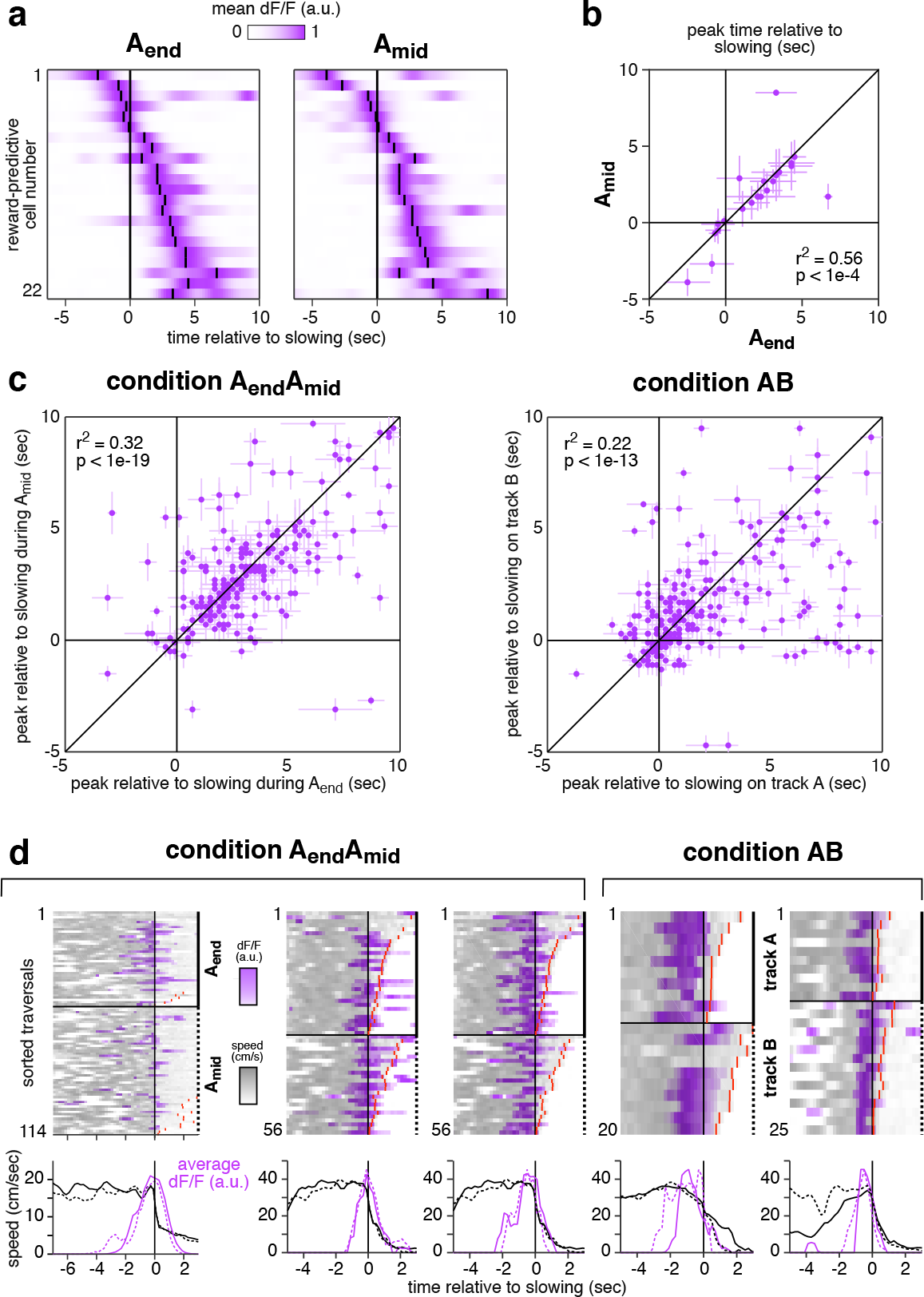
Reward-predictive cells formed a consistent sequence that began prior to reward anticipation behavior. **a.** Mean activity of 22 simultaneously-recorded reward-predictive cells from CA1 shown in the same order for A_end_ (left) and A_mid_ (right). Small black lines indicate time of peak activity. Cells were selected for having COM locations within 50 cm before reward and being active on at least 20 trials in the 100 cm before reward. Time courses were filtered with a Gaussian kernel (s.d. 0.1 sec). **b.** Time of peak activity relative to slowing for the same cells as in a. Bars indicate width at half max of unfiltered trace. **c.** Time of peak activity relative to slowing for all reward-associated cells recorded during condition A_end_A_mid_ (218 cells, 6 mice) and condition AB (243 cells, 5 mice). Bars indicate width at half max of unfiltered trace. **d.** Five reward-predictive cells active early in their respective sequences. In each case, fluorescence increased 1-2 seconds before speed decreased. Cells were recorded in four mice, each column a cell. Cells in column 2 and 3 were recorded simultaneously. The cell in column 4 was from subiculum, and others were from CA1. Top: speed and activity on single traversals, same conventions as in Figure 4f. Activity on each trial was normalized to have a maximum of 1. Red lines indicate the time of reward delivery when it occurred early enough to be within plot bounds. Bottom: average across trials of activity (80th percentile) and speed (mean).

Activations in approximately the same order were also found for the full populations of reward predictive cells in both conditions (Figure 4c), and also when considering CA1 and subiculum separately (not shown). Such similar sequential activation, regardless of location or environment, showed that even individual reward-predictive cells seemed to be highly specialized.

### 2.5. Reward-predictive cell sequences did not encode reward anticipation behaviors

While the previous findings were consistent with the interpretation that reward-associated cells provided a reliable marker of reward location that might support goal-directed navigation, they also raised a very different possibility: that reward-predictive cells simply encoded the motor or sensory events that coincided with reward anticipation. This seemed unlikely, given that not all reward-predictive cells were correlated with slowing, and that different cognitive demands resulted in different fractions of the population exhibiting correlation.

Nevertheless, this possibility was tested in four control analyses that compared reward-predictive cell activity to behavior in several respects. First, the relative timing of activity and behavior was compared. If reward-predictive cell sequences were triggered by slowing per se, they would always start after speed began to decrease. Contrary to this prediction, in several preparations the sequence of reward-predictive cells became active prior to any detectable reduction in speed (Figure 4d).

To show that reward-predictive cells did not encode events preceding a decrease in speed, such as changing gait or premotor planning, reward approach was compared to other instances in which mice slowed down. While running between reward locations during condition A_end_A_mid_, mice occasionally slowed, stopped, then resumed running. These brief rest events were initiated at locations throughout the track and were almost never accompanied by licking (Supplemental Figure 5), suggesting they were unrelated to reward anticipation. At the onset of rest events, reward-predictive cell activity was indistinguishable from the baseline activity during running (0.9 ± 0.4 %ΔF/F, mean ± std. err. vs 1.12 ± 0.03, p = 0.57, student’s t-test), and far below the average activity observed when mice slowed prior to reward (7.1 ± 0.8). In the first 1 second after slowing, activity during rest events remained indistinguishable from baseline (1.3 ± 0.03, p = 0.65), while during prereward walking bouts it increased even further (10.4 ± 0.7). These comparisons demonstrated that reward-predictive cells did not encode events associated with slowing, since their activity remained at baseline levels when slowing was unrelated to reward anticipation.

It was also shown that reward-predictive cell activity could not be explained as a simple function of licking. Comparing the three-second intervals just before and after reward delivery, the lick rate increased more than four-fold (1.01 ± 0.01 Hz pre vs 4.73 ± 0.01 post), while the fluorescence of reward-predictive cells fell by nearly half (11.5 ± 0.1 %Δ*F/F* pre vs 5.99 ± 0.08 post).

Finally, it was shown that reward-predictive cells were not responding to the full constellation of anticipation behaviors, i.e. slowing down, walking at a low speed for several seconds, and simultaneously licking. This possibility was tested using a natural control: “error” trials.

During condition A_end_A_mid_, mice frequently slowed, walked, and licked prior to reward. But sometimes they exhibited these behaviors at other locations, especially before the rewarded location of the non-current context (e.g. walking before 166cm when the reward was delivered at 366cm, Supplemental Figure 6). These “incorrect” walking bouts were defined as those overlapping the non-current reward location (see Methods), and they were accompanied by significantly more licking than walking bouts that did not overlap a rewarded location (2.12 vs 0.85 licks/bout, p < 1e-10, student’s t-test), showing that mice had an expectation of reward. If reward-predictive cells encoded the behavioral events associated with reward anticipation, they should exhibit the same level of activity regardless of where anticipation took place, and in particular they should be equally active when walking before the current or non-current reward location.

Figure 5a shows two example walking bouts from the same session, one before the current reward location, and one spanning the non-current reward location. In both cases, the mouse suddenly slowed down, then walked for several seconds while licking at a rate of approximately 1 Hz. Despite the similarity of anticipation behaviors, reward-predictive cells were much more active when approaching the current reward site than the non-current reward site.

**Figure 5.**
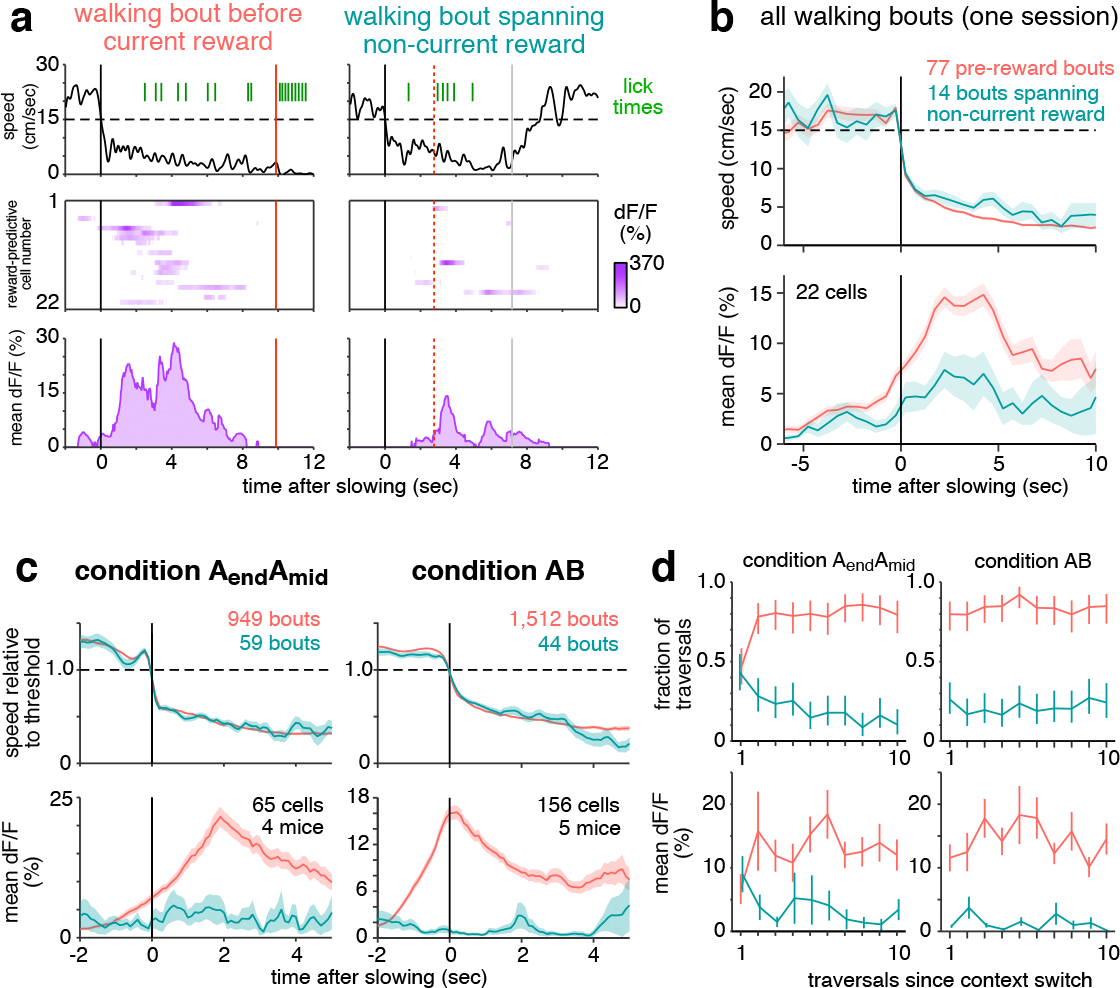
Reward-predictive cell activity can not be explained by reward anticipation behavior. **a-b.** Activity is shown for the same 22 simultaneously-recorded reward-predictive cells depicted in Figure 4a. **a.** Instantaneous movement speed (top), activity of each reward-predictive cell plotted in the same order as in Figure 4a (middle), and population mean activity (bottom) during two walking bouts, preceding the current (left) or non-current reward location (right). Black lines indicate onset of walking, red lines indicate reward, and gray lines indicate the end of the unrewarded walking bout. **b.** Top: Movement speed averaged over all walking bouts from this session, excluding the first three traversals of each block, grouped by whether they preceded current (pink) or non-current (blue-green) reward. Bottom: Simultaneous activity of reward-predictive cells. Single trial traces were averaged in half-second chunks before combining across trials, bands show standard error of the mean across trials. **c.** Average speed relative to slowing threshold (top panels) and average activity of reward-predictive cells (bottom). Only includes sessions in which reward-predictive cells were recorded. A subset of bouts was manually chosen (see Methods) to maximize the similarity of average speed; for all bouts, see Supplemental Figure 7. Condition A_end_A_mid_ only includes data from day 7 of training or later to ensure mice were familiar with the reward delivery paradigm. **d**. Comparison of how quickly slowing behavior and reward-predictive cell activity adapt to a new context. Condition AB only includes data from track A. Top: fraction of traversals in which mice exhibited a pre-reward walking bout (pink) or an unrewarded walking spanning the non-current reward location (blue-green). Error bars indicate 95% confidence interval. Bottom: mean fluorescence of reward-predictive cells in the first 5 seconds after slowing onset, error bars show standard error of the mean.

These examples are representative of the entire session (Figure 5b). Average movement speeds were virtually identical in the two categories, yet reward-predictive cell activity was more than two times greater when walking before the current reward. An even greater difference in activity was observed when considering the full population of reward-predictive cells recorded from mice in both conditions (Figure 5c). Several control analyses verified that differential activity could not be ascribed to a difference in the average lick rate, the overall level of hippocampal activity, or selection bias introduced when classifying reward-predictive cells (Supplemental Figures 7 and 8).

This comparison demonstrated that reward-predictive cells did not encode the behavioral events that typically preceded reward. Instead of producing a stereotyped response to all instances of reward anticipation, their activity was strongly modulated by the particular circumstances in which anticipation took place. This suggested that they encoded a cognitive variable that reflected the internal state, one that apparently differed when the mouse was walking at the current or noncurrent reward location.

Because the previous analyses averaged across trials and excluded the first three traversals of each block, a separate analysis was performed to identify how rapidly reward-predictive cells remapped after the context switched (Figure 5d). During condition A_end_A_mid_, activity shifted after one or two exposures to the new reward location, whereas during condition AB the change was immediate. These time courses were similar to the speed at which reward anticipation shifted to the new location, revealing one more respect in which reward-predictive cell activity was aligned to changes in behavior.

## 3. Discussion

We have described a novel population of neurons in two major hippocampal output structures, CA1 and the subiculum. Reward-associated cells exhibited activity fields that did not deviate from the reward location, and these cells entirely accounted for the excess density of fields near reward. Their pattern of remapping cleanly distinguished them from simultaneously-recorded place cells, both when reward was shifted within one environment and most place fields remained stable, as well as across environments when place cells remapped to random locations. During these manipulations, the population of reward-associated cells did not mix with place cells, indicating they composed a distinct subgroup. Examining the single-trial timing of reward-predictive cells showed that they were often precisely aligned to reward anticipation, whereas place cells tended to represent spatial locations independently of behavior. These findings demonstrate an unexpected degree of consistency in the hippocampal representation of reward, and the observed properties of reward-associated cells suggest they might play an important role in navigation.

Reward-associated cells appear to be the experimental confirmation of a cell class hypothesized more than two decades ago [6]. Burgess and O’Keefe proposed a dedicated class of “goal cells” that would serve as an anchor point for the hippocampus to devise goal-directed trajectories. More recently, other models of navigation have been developed in which reward-associated cells could serve a critical function. For example, animals might explore possible routes in advance of movement [36, 20], in which case activating reward-associated cells would indicate a successful route. Other observations have suggested the computation proceeds in reverse, from the goal to the current location [1], meaning reward-associated cells could provide a seed for this chain of activations. In the separate framework of reinforcement learning, reward-predictive cells might confirm predictions of the successor representation theory [11, 49], in that they signal an impending reward. While diverse in their algorithms, these models illustrate the importance of reward-associated fields being carried by the same cells: consistency enables other circuits to reliably identify reward location, regardless of environment or context.

Though reward-associated cells were not characterized anatomically, it is possible that they project to a specific external target, such as nucleus accumbens. This would be consistent with recent observations in ventral CA1 that neurons projecting to nucleus accumbens are more likely to be active near reward than neurons with other projections [8]. If reward-associated cell axons did reach the ventral striatum, their sequential activation could conceivably [16] underlie the ramping spike rate that precedes reward [5], and might even contribute to reward prediction error signals of dopaminergic neurons [46].

Reward-predictive cell sequences were often precisely aligned with, and sometimes even preceded, anticipation behaviors. This might have reflected reward-predictive cells making a direct contribution to reward anticipation, or they might have received an “efferent copy” of a prediction signal generated elsewhere. It is intriguing that more cells were correlated with behavior during anticipation that required short term memory, which is generally associated with hippocampus-dependent tasks [22, 44], and less prevalent when anticipation could have based entirely on immediate cue association, which typically does not rely on the hippocampus [40].

An important open question is why reward-predictive cells were more active when mice anticipated reward correctly than incorrectly. The amplitude of activity might have encoded the level of subjective confidence that reward was nearby, possibly implying a link to orbitofrontal cortex neurons that seem to encode value [45]. Alternatively, the differing amplitudes might reflect the existence of multiple reward prediction systems [10], with the hippocampus contributing a prediction in some, but not all, bouts of reward anticipation.

Given their unique physiology, it is striking that, to our knowledge, reward-associated cells have not been described previously, despite decades of research on hippocampal activity in the context of reward. One possible explanation is that their sparsity (1-5% of recorded neurons) made them difficult to detect, especially prior to the advent of large-scale recording technologies. Nevertheless, they might be related to CA1 neurons in bats that seem to encode a vectorial representation of the goal [42].

It is also possible that reward-associated cell activity is only elicited during specific circumstances. Comparing across several studies with different task structures [23, 15, 9, 43], two task features seem required to observe reward-associated cells: no explicit cues for reward, and reward locations frequently shifting to different parts of the environment. These findings suggest that reward-associated cells are not a general feature of hippocampal encoding, but instead are only engaged in the context of particular cognitive demands.

Whatever conditions might elicit the activity of reward-associated cells, it is clear that they encode a variable of central importance for goal-directed navigation, and they endow hippocampal maps with a consistency that was not previously appreciated. In addition, they provide a novel target for studying the hippocampal contribution to reward memory. Further studies will be required to determine whether reward-associated cells relate to the encoding, storage, or recall of reward locations, and how they might interface with other brain areas to support navigation.

---

**Supplemental Figure. 1.**
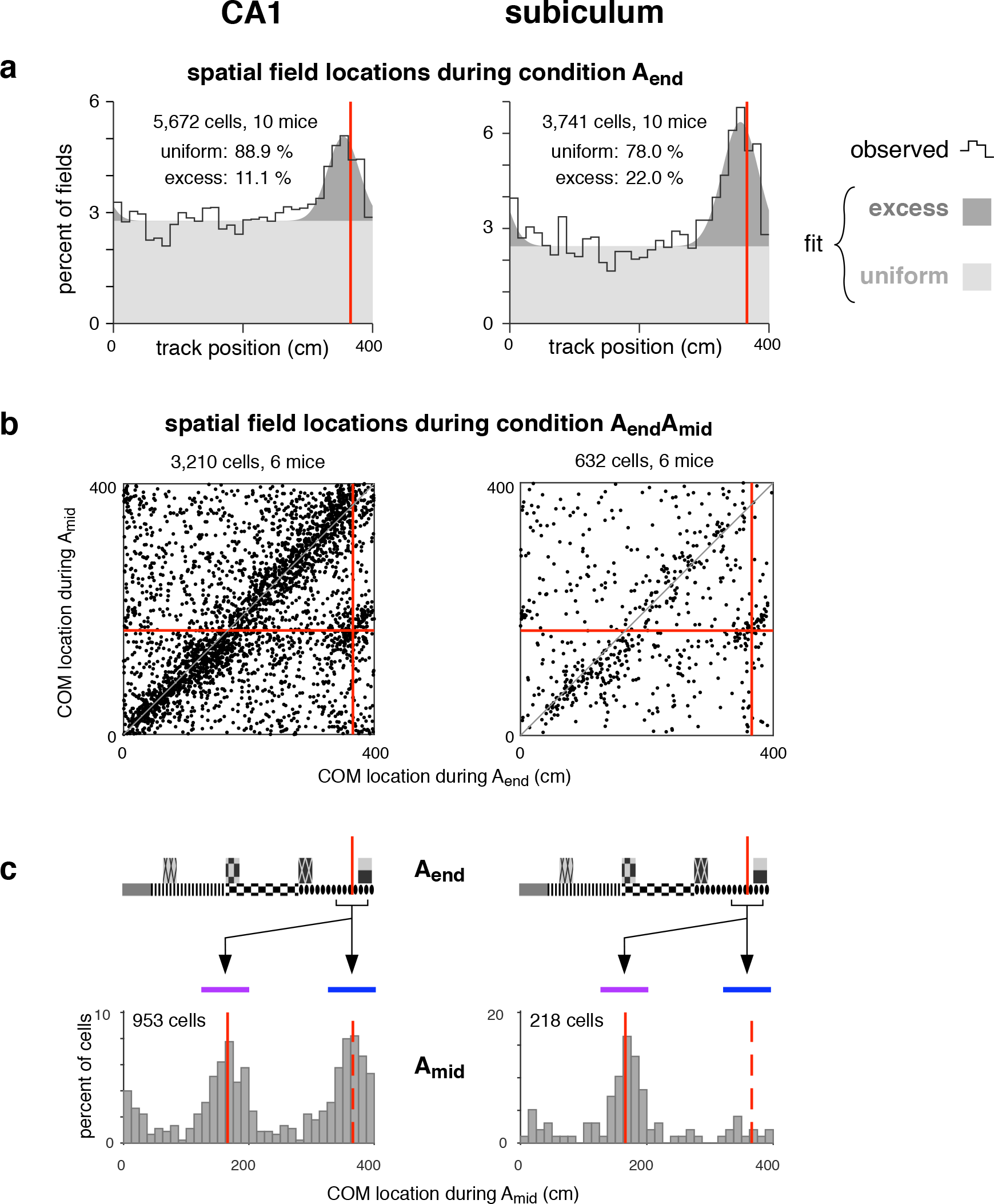
Reward-associated cells were found in both CA1 and subiculum. **a-c.** Analyses were performed separately for CA1 (left panels) and subiculum (right panels). **a.** Density of spatial field COMs. Black line shows observed density, gray patches show density of a fitted mixture distribution consisting of a uniform distribution (light gray) and a Gaussian distribution (dark gray, mean 355 cm, s.d. 24 cm for CA1, mean 356 cm, s.d. 28 cm for subiculum). **b.** COM locations of cells with a spatial field during both A_end_ and A_mid_. Same graphical conventions as in Figure 1g. **c.** COM location during A_mid_ for cells with a COM located within 25 cm (before or after) reward during A_end_. Same graphical conventions as in Figure 1f. Although very few place cells in the subiculum were located near 366 cm, there was still a population of reward-associated cells whose fields remapped to be consistently active near reward.

**Supplemental Figure. 2.**
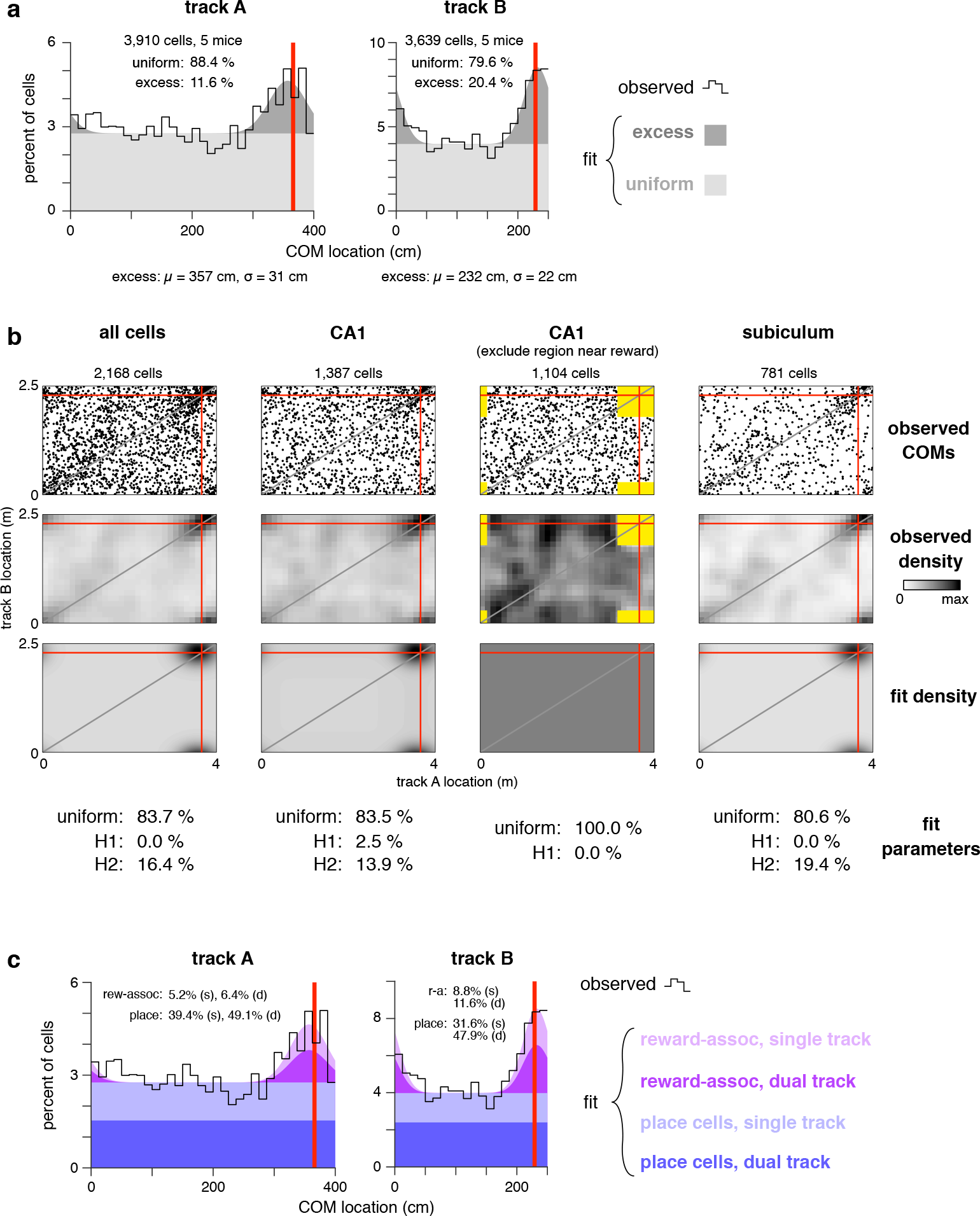
Analysis of COM density in condition AB. **a-c.** Red lines indicate reward location. **a.** During condition AB, COMs were located at excess density near reward on both track A (left) and track B (right). Black outlines show observed COM density, and solid patches show fit density of uniform distribution (light gray) and the excess density near reward modeled as a Gaussian (dark gray). Inset numbers show fit coefficients of the mixture distribution. Numbers at bottom show parameters of Gaussian fit. **b.** Among cells with a spatial field on both tracks, the excess density was entirely explained by H2 rather than H1 (see Results and Methods). Each column shows analysis of COMs from a different cell group. First row: observed COM locations. Second row: observed COM density, binned in 12.5 cm bins, smoothed with a Gaussian kernel (15 cm radius). Third row: Density of fit mixture distribution. Fourth row: fit coefficients of mixture distribution. The fit to all CA1 neurons (second column) seemed to indicate some cells remapped according to H1 (2.5%). However, this was an artifact caused by the density near the reward being poorly fit by a Gaussian. After excluding the region within 50 cm of both rewards (yellow patches in third column), there was no contribution from H1 (0.0%). **c.** The observed distribution of COM locations (black outline) was modeled as a collection of place cells and reward-associated cells (colored patches) based on the results of the H1-H2 fits (see Methods). The excess COM density near reward was entirely composed of reward-associated cells. The fitted model was further decomposed into populations of cells with a spatial field on one track or on both tracks.

**Supplemental Figure. 3.**
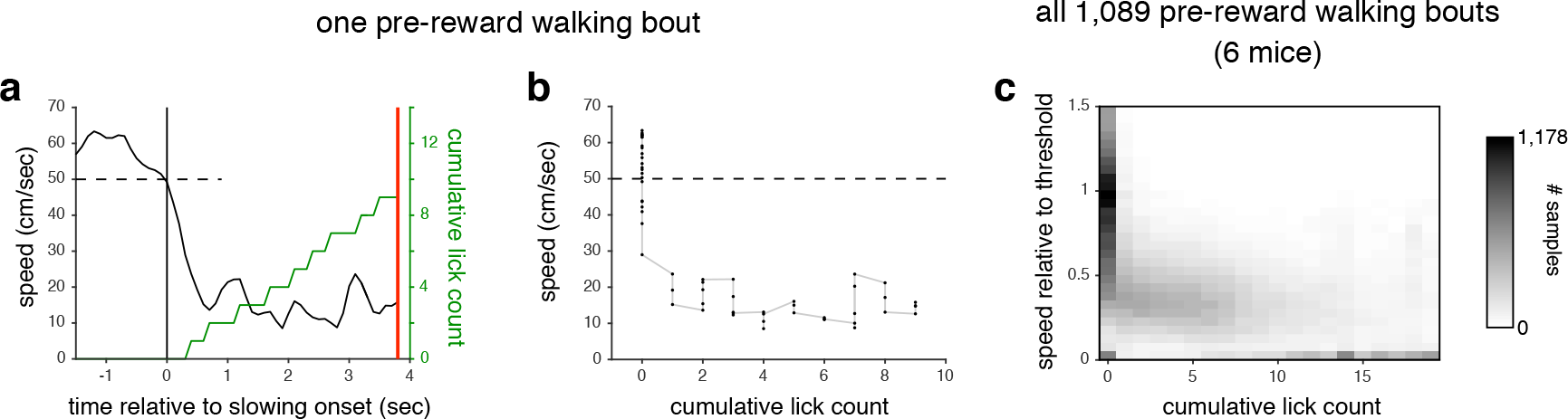
During reward approach, slowing preceded walking. **a.** Speed (black) and cumulative lick rate (green) sampled at 10 Hz for a representative pre-reward walking bout. Dashed line indicates speed threshold. Red line indicates reward delivery. **b.** Scatter plot of speed vs cumulative lick rate (same data as panel a). **c.** Density of speed vs cumulative lick rate for all pre-reward walking bouts with at least 5 licks during condition A_end_A_mid_. Values were collected in the time interval from 2 seconds prior to slowing until reward delivery. In nearly every case, speed was significantly reduced prior to the first lick, showing that slowing preceded licking.

**Supplemental Figure. 4.**
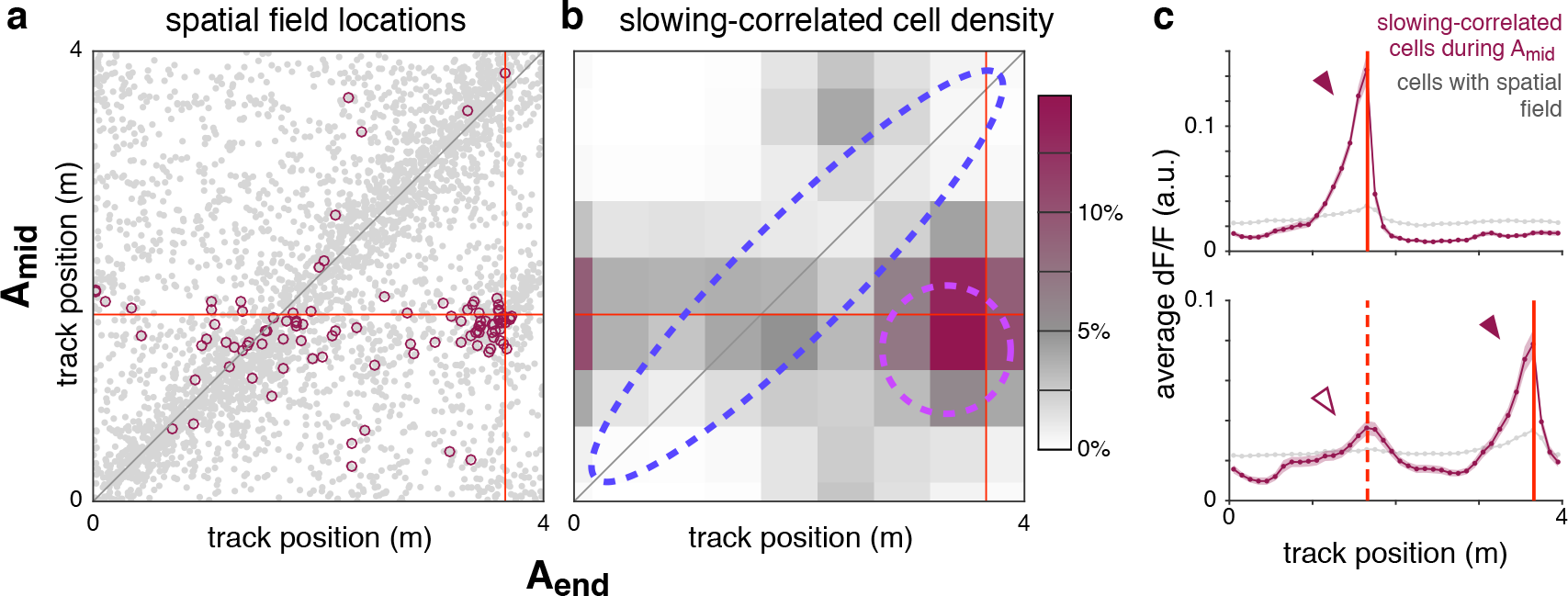
During A_mid_, only reward-predictive cells are slowing-correlated above chance levels. Same analysis as main Figure 3i,j applied to cells that were slowing-correlated during blocks of context A_mid_. **a.** COM locations of cells with spatially-modulated fields in both A_end_ and A_mid_ (gray points, same data as Figure 1g). Cells are highlighted (maroon circles, 103 cells) if they were slowing-correlated during A_mid_ (see Results for definition). **b.** Lower bound of estimated density of slowing-correlated cells (see Methods). Dashed lines indicate approximate boundaries of regions used to define reward-predictive cells (purple) and place cells that were stable across conditions (blue). **c.** Upper: average fluorescence activity of 142 slowing-correlated cells (6 mice, maroon trace) during A_mid_ blocks, and all spatially-modulated cells recorded simultaneously (6,935 cells, gray trace), plotted as a function of track position. Red line indicates A_mid_ reward location. As expected, slowing-correlated cells exhibited increased activity just prior to the reward (arrowhead). Lower: same averaging procedure applied to same cells, but during the interleaved A_end_ blocks from the same sessions. Red lines indicate reward location for A_end_ (solid) and, for comparison, A_mid_ (dashed). Bands indicate standard error of the mean. Despite some residual above-baseline fluorescence at the A_mid_ reward site (hollow arrowhead), the majority of slowing-correlated cell activity shifted to the A_end_ reward site (solid arrowhead), consistent with few if any place cells being slowing correlated.

**Supplemental Figure. 5.**
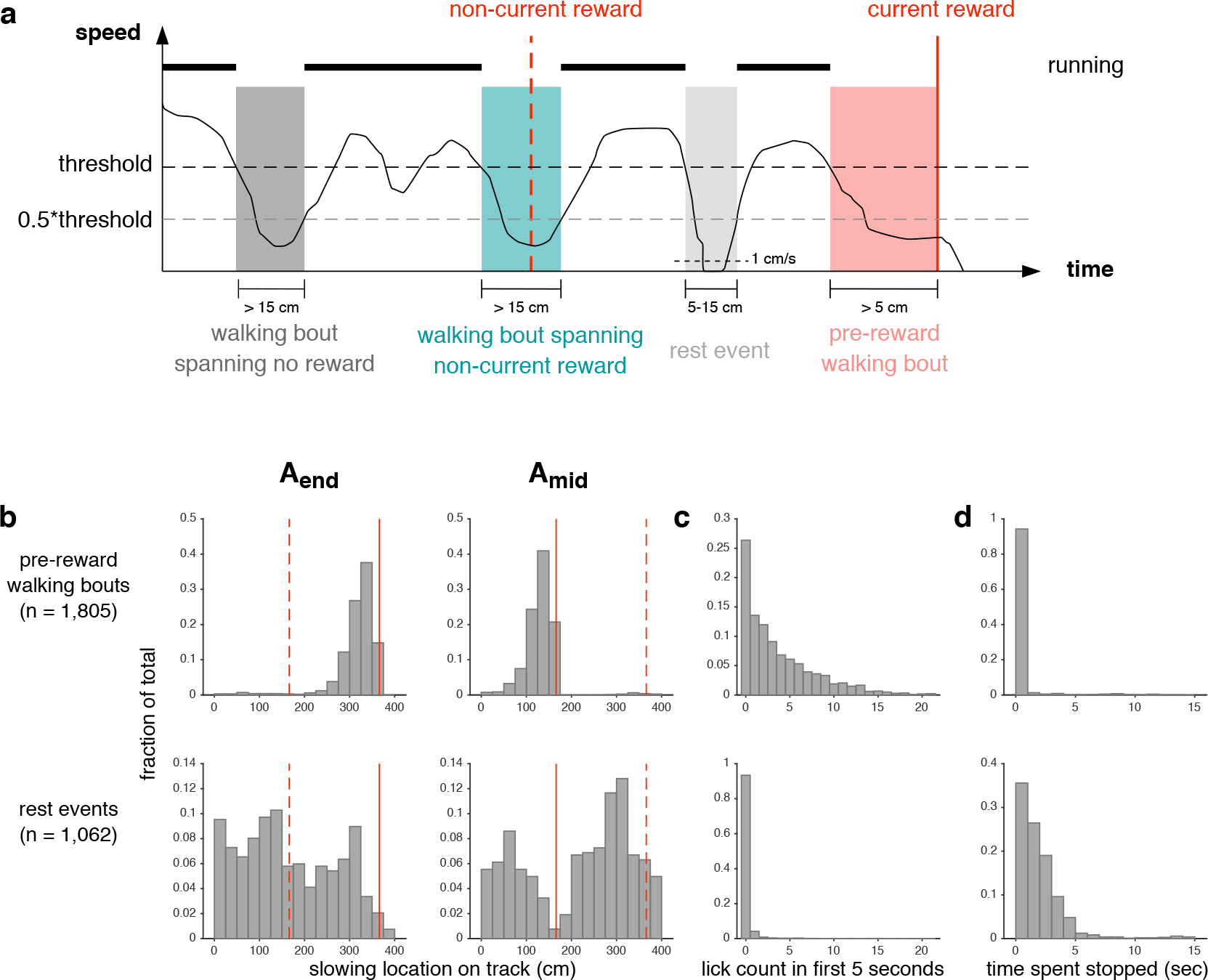
Brief rest events do not reflect anticipation of reward. **a.** Schematic illustrating how changes in movement speed (black trace) are used to define walking and rest events (colored patches). For detailed definitions, see Methods. **b.** Locations where pre-reward walking bouts (top) or rest events (bottom) were initiated, plotted separately for blocks of A_end_ (left) or A_mid_ (right). To ensure mice were familiar with the structure of reward delivery and did not slow because they expected reward at other locations, data was only included from session 7 or later of training on condition A_end_A_mid_, and walking bouts from the first three traversals of each block were excluded. **c.** Number of licks during the first 5 seconds after the walking bout or rest event began. **d.** Overall duration in which mice were stopped (i.e. speed was slower than 1 cm/sec).

**Supplemental Figure. 6.**
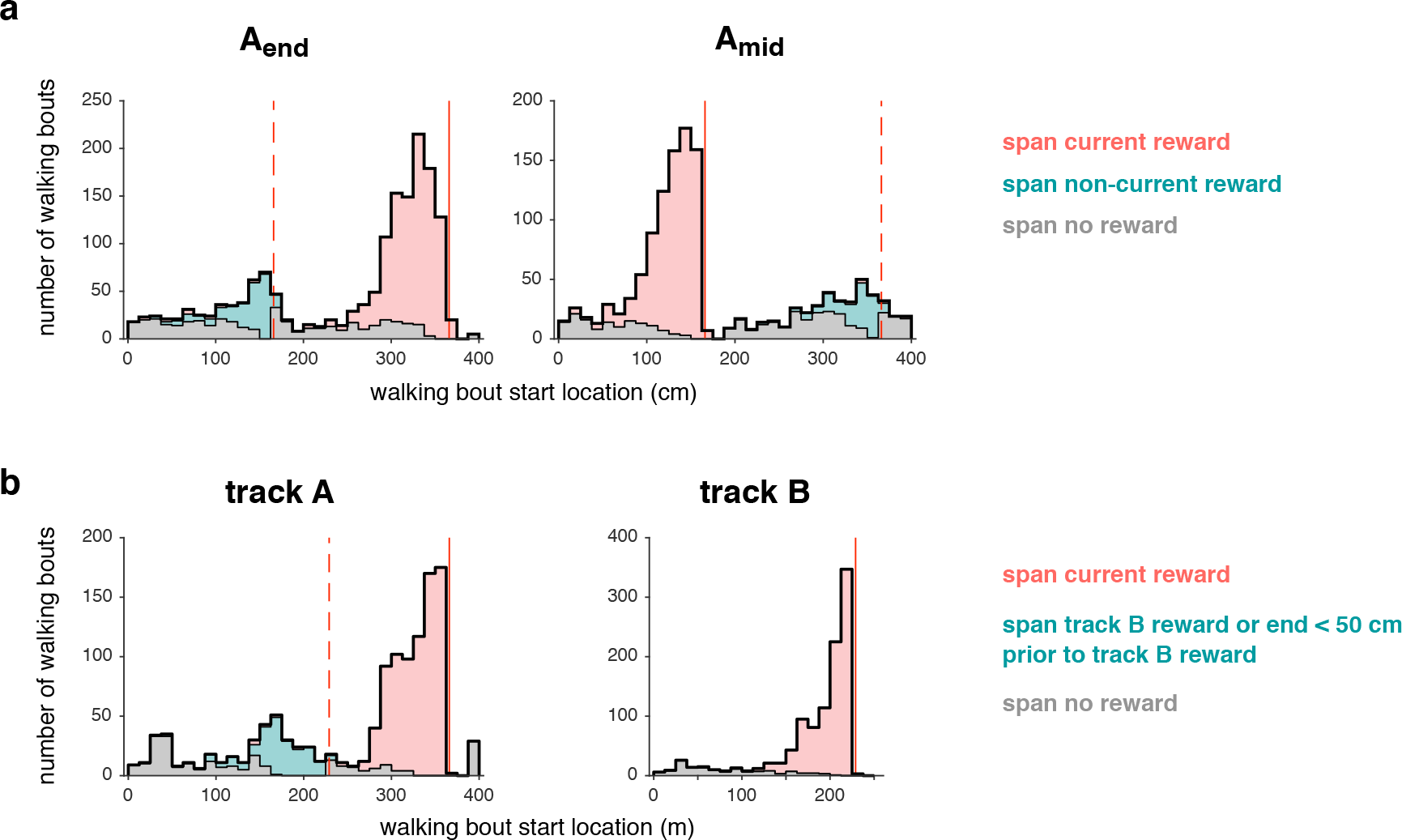
Mice bouts were clustered near the current and non-current reward locations. The first three traversals of each block were excluded, and for condition A_end_A_mid_ data are only shown for session 7 or later. **a.** Starting locations for all walking bouts during condition A_end_A_mid_, grouped by whether they spanned the current reward (pink), non-current reward (blue-green), or no reward (gray). **b.** Equivalent analysis for condition AB. Since the two rewards were delivered on different tracks, there was not an obvious definition of the “non-current reward location”. Nevertheless, mice running on track A exhibited an increased frequency of slowing in the 150-200 cm range, presumably since this was close to 229 cm (location of track B reward). To ensure these walking bouts were considered related to the non-current reward location, the definition was expanded to include bouts that spanned any point in the 50 cm preceding 229 cm.

**Supplemental Figure. 7.**
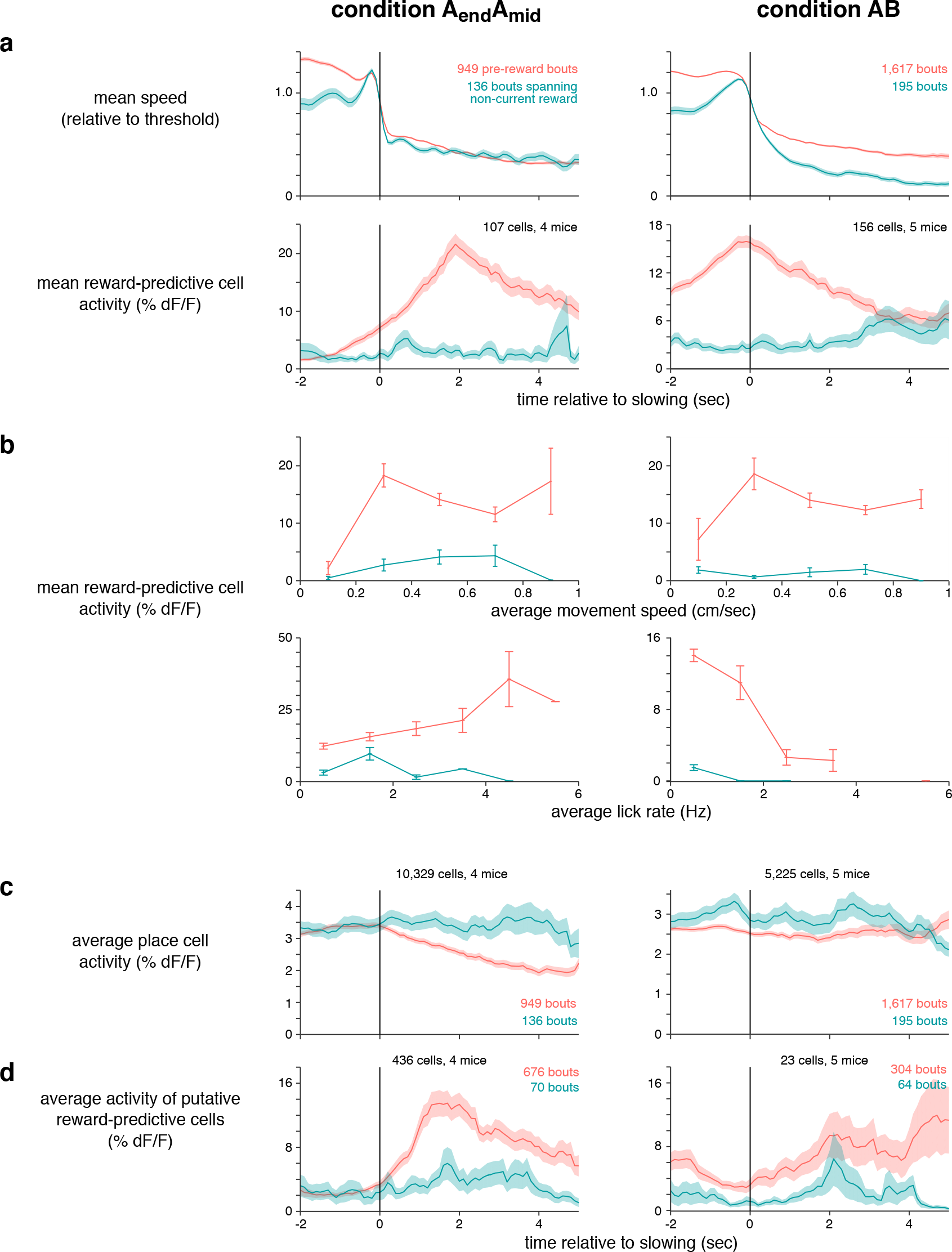
Modulation of reward predictive-cell activity could not be explained by speed, licking, overall activity, or selection bias. All plots compare activity or speed when mice walked before the current reward location (pink) or non-current reward location (blue-green). Error bars or bands indicate standard error of the mean when averaging across bouts. Activity of cells was first averaged within each bout, meaning error bars for activity overestimate the true uncertainty. For condition A_end_A_mid_, data are only shown for day 7 of training or later and the first three traversals of each block were excluded. Speed data only includes sessions in which reward-predictive cells were recorded. **a.** Mean speed (top) and reward-predictive cell activity (bottom) of slowing bouts as a function of time relative to slowing. **b.** Activity of reward-predictive cells when bouts were subdivided based on movement speed (top) or lick rate in the first 5 seconds after slowing (bottom). **c.** Activity of all cells that exhibited a spatially-modulated field in at least one context and were not classified as reward-predictive cells. Only includes sessions in which reward-predictive cells were also recorded. Though overall activity levels differed slightly between approaching current or non-current reward, they could not account for the difference in activity of reward-predictive cells. **d.** Activity of putative reward-predictive cells, analyzed to avoid selection bias (see Methods). In condition A_end_A_mid_, activity was much greater when approaching the current reward, confirming that the effect shown in Figure 5 could not be attributed to selection bias. In condition AB, the putative reward-predictive cells might have included many place cells that were spuriously slowing-correlated on track B, and thus remapped randomly on track A, contributing a uniform positive offset in the pink trace.

**Supplemental Figure. 8.**
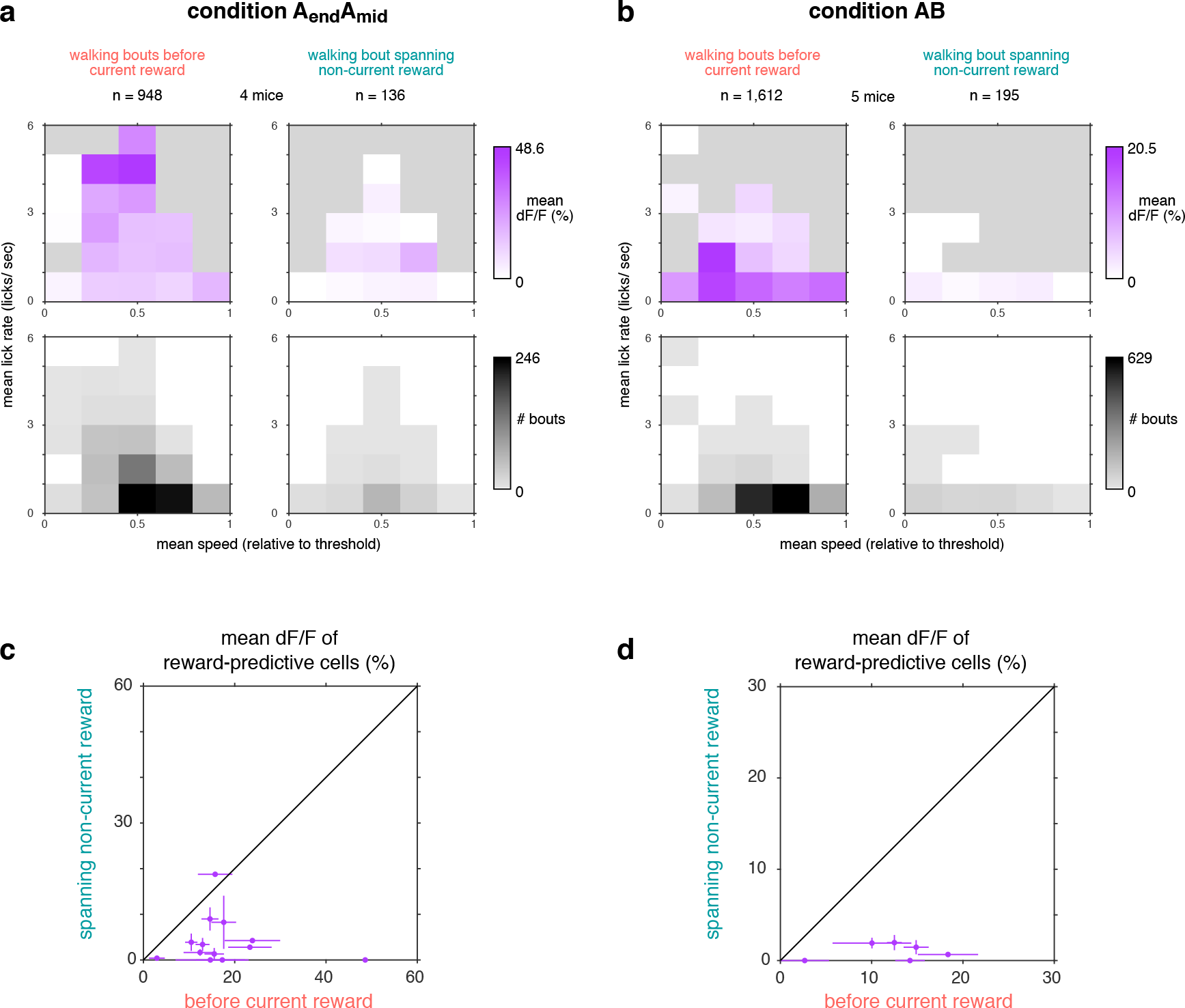
Modulation of reward predictive-cell activity could not be explained by the conjunction of speed and lick rate. **a.** Each bout was categorized based on the average lick rate (y-axis) and average speed (x-axis) in the first 5 seconds. Top panels: Average reward-predictive cell activity during bouts in each category. Bottom panels: number of bouts in each category. Separate plots are shown for pre-reward walking bouts (left) and walking bouts spanning the non-current reward (right). Data are only shown for day 7 of training or later and the first three traversals of each block were excluded. **b.** Equivalent analysis for condition AB, except that all training days and traversals were included. **c.** Scatter plot of reward predictive-cell activity during pre-reward walking bouts (x-axis) and walking bouts spanning the non-current reward (y-axis). Error bars indicate standard error of the mean. Each point is the average of bouts from one category (i.e. a single bin from panel a), allowing a direct comparison of reward-predictive cell activity when controlling for the conjunction of lick rate and walking speed. In nearly every case, activity was greater when walking before the current reward than the non-current reward, confirming the result of Figure 5. **d.** Equivalent analysis of data in panel b.

## 4. Methods

### 4.1. Surgery

All experiments were performed in compliance with the Guide for the Care and Use of Laboratory Animals (http://www.nap.edu/readingroom/books/labrats/). Specific protocols were approved by the Princeton University Institutional Animal Care and Use Committee.

Transgenic mice expressing GCaMP3 [39] (C57BL/6J-Tg(Thy1-GCaMP3)GP2.11Dkim/J, Jackson Labs strain 028277) were used to obtain chronic expression of calcium indicator. Optical access to the hippocampus was obtained as described previously [13]. A small volume of cortex overlying the hippocampus was aspirated and a metal cannula with a coverglass attached to the bottom was implanted. A thin layer of Kwik-Sil (WPI) provided a stabilizing interface between the glass and the brain. The craniotomy was centered at the border of CA1 and subiculum (1.8 mm from the midline, 3 mm poster to bregma) so that both regions could be imaged in a single window, though not simultaneously. During the same surgery a metal head plate was affixed to the skull to provide an interface for head fixation.

Mice and their littermates were housed together until surgical implantation of the optical window. After surgery, mice were individually housed. Mice were housed in transparent cages on a reverse light cycle, with behavioral sessions occurring during the dark phase.

### 4.2. Behavioral training

At the time of surgery, mice were aged 7 to 15 weeks. After recovering for at least 7 days, water intake was restricted to 1 to 2 ml of water per day and was adjusted within this range based on body weight, toleration of water restriction, and behavioral performance. After several days of water restriction, mice began training in the virtual environment, typically one session per day and 5-7 days per week.

The virtual reality enclosure was similar to that described previously [13, 14]. Briefly, head-fixed mice ran on a styrofoam wheel (diameter 15.2 cm) whose motion advanced their position on a virtual linear track, and an image of the virtual environment was projected onto a surrounding toroidal screen. The virtual environment was created and displayed using the VirMEn engine [4]. To mitigate the risk of stray light interfering with imaging of neural activity, only the blue channel of the projector was used, and a blue filter was placed in front of the projector.

Condition A_end_: The virtual track was 4 meters long, with a variety of wall textures and towers that served to provide a unique visual scene at each point on the track. Textures and tower locations were chosen to replicate as closely as possible a track used in a previous study [14]. When mice reached a point just before the end (366 cm), a small water reward (4 uL) was delivered via a metal tube that was always present near the mouth. The reward location in the virtual environment was unmarked, insofar as visual features at that location were no more salient than at other points on the track. After running to the end of the track, mice were teleported back to the beginning. To avoid visual discontinuity, a copy of the environment was visible after the end of the track.

After each reward was delivered, the small droplet of water remained at the end of the tube and was available for consumption indefinitely. When mice licked the reward tube, regardless of whether water was available, each lick was detected using an electrical circuit that measured the resistance between the mouse’s head plate and reward tube. The resistance was sampled at 10 kHz, and licks appeared as brief (10-20 msec) square pulses. Before identifying lick onset times, a Haar wavelet reconstruction was performed to reduce electrical noise. In a few cases, electrical noise was large enough to interfere with lick detection, and these datasets were excluded from analyses that involved licking.

After at least 5 sessions of training on condition A_end_, mice were exposed to a new reward delivery paradigm, either condition A_end_A_mid_ or condition AB.

Condition A_end_A_mid_: The reward location alternated block-wise between 366 cm and 166 cm (condition A_end_A_mid_, Figure 1d). Within each block, the reward was delivered at either 366 cm (A_end_) or 166 cm (A_mid_). Each session began with a block of context A_end_. Block transitions occurred seamlessly, with no explicit cue indicating that the reward location had changed. The reward locations were not explicitly marked, and there were no visual features common to the two reward locations that distinguished them from other parts of the track.

Condition AB: Within each block, mice either traversed track A (400 cm, reward at 366 cm) or track B (250 cm, reward at 229 cm). The two tracks had no common visual textures. Block changes took place during teleportation at the end of the track, creating a brief visual discontinuity. Each session began with a block of track A.

Block durations: When a new block began, two criteria were chosen to determine when to switch to the next block. One criterion was a time interval, typically chosen randomly between 5 and 15 minutes, and the other criterion was a number of traversals, typically chosen randomly in the range 10 to 20. When either the amount of time or the number of traversals had been reached, the context changed at the next teleport and a new block began. Across all sessions and mice, the average block duration was 8.4 ± 5.9 minutes (mean ± s.d.) and the average number of rewards was 18.7 ± 13.4.

Imaging windows were implanted in a total of 24 mice. Of these, 3 mice were excluded because of poor imaging quality, 8 were excluded because of poor behavior in condition A_end_ (typically earning less than 1 reward per minute), 1 died unexpectedly, and 12 were used in the study. Separate cohorts of mice were used for condition AB (5 mice) and condition A_end_A_mid_ (7 mice), though one mouse whose data was used for condition A_end_A_mid_ had previously been exposed to 10 sessions of condition AB (data from the condition AB sessions was not used due to a problem with experimental records).

### 4.3. Optical recording of activity

While mice interacted with the virtual environment, two-photon laser scanning microscopy was used to identify changes in fluorescence of the calcium indicator GCaMP3 caused by neural activity. In most experiments (see exception below), the two-photon microscope was the same as described previously [13]. Typical fields of view measured 100 by 200 um, and were acquired at 11-15 Hz.

In CA1, approximately half of pyramidal neurons were labeled, specifically those located in the dorsal half of the pyramidal layer. In subiculum, approximately three quarters of cells were labeled, with labeled cells distributed throughout all depths.

To examine the population activity of many simultaneously-recorded cells during a single session, additional data was obtained from CA1 in one mouse with modified experimental parameters: individual blocks and sessions lasted longer, and a larger field of view was imaged (500 × 500 um) at a faster scan rate (30 Hz). To obtain a larger field of view, a modified version of the two-photon microscope was used, similar to a design described previously [26]. Compared to the microscope used in other experiments, the most significant change was the incorporation of resonant galvanometer scan mirrors.

This mouse (EM7) was trained on condition A_end_A_mid_, and data from only the longest session (number 12, 114 traversals) was used here. This session provided example data for several figure panels (3d-h, 4a,b,d, and 5a-b).

However, data from this mouse was not used in the population analyses. The same field of view was imaged on each day of behavioral training, but no attempt was made to track single cells. When all recorded cells from this mouse were pooled over time, reward-associated cells were observed, confirming suitability of the data as representative of the other mice described in this study. Nevertheless, if this pooled data had been included in the population analyses, it would have introduced many unidentified duplicate cells, potentially biasing the results, and not being compatible with some statistical tests.

### 4.4. Identification of cell activity

All analyses were performed using custom software in MATLAB (Mathworks).

Motion correction of recorded movies was performed using an algorithm described previously [13]. Cell shapes and fluorescence transient waveforms were identified using a modified version of an existing algorithm [30]. The principal modification was in using a different normalization procedure: instead of dividing each frame by the baseline, each frame was divided by the square root of the baseline to yield approximately the same resting noise level in all pixels. This normalization was used only to identify cell shapes, but not for extracting time courses (see below).

In each movie, the algorithm typically identified 30-150 active spatial components (each referred to as a “cell”). All cells were kept for subsequent analyses, with no attempt to distinguish somata from processes. Time courses were computed as follows. For each pixel in each frame, the baseline was computed by taking the 8th percentile of values in that pixel in a rolling window of 500 frames. In each frame, the activity amplitude of all cells was computed by performing a least-squares fit of the cell shapes to the baseline-subtracted frame, yielding the fractional change in fluorescence, or Δ*F/F*. For each cell, the time course was median-filtered (length 3) and thresholded by zeroing time points that were not part of a significant transient at a 2% false positive rate [13].

In some cases, the motion-corrected movie contained a small amount of residual displacement in the Z axis, typically about 1 micron. Though small, this displacement could produce apparent changes in the fluorescence of up to 50%. Because Z displacement was uniform over the entire image, its value could be readily measured at single-frame time resolution, yielding an estimated Z displacement time course. In the time course of each cell, the amplitude of the Z displacement time course was fitted and subtracted before the filtering and thresholding steps described above. This prevented artifactual changes in fluorescence from contaminating true transients.

In the dataset that employed resonant scan mirrors to obtain a wider field of view, the above methods could not be applied. The field of view was so large that motion offsets were not consistent throughout the image (e.g. the top of the image was displaced right while the bottom was displaced left), which necessitated a more complex motion correction procedure.

First, whole-frame correction was applied separately to each chunk of 1000 frames using the standard algorithm. To correct for residual motion within each frame, the corrected movie was divided into 5 spatial blocks, each of which spanned the entire horizontal extent of the image. Vertically, blocks were evenly sized and spaced, and adjacent blocks overlapped by 50%. In each of the 5 blocks, motion was identified using the standard algorithm, and these offsets were stored for subsequent correction.

For cell finding, the imaged area was divided into 36 spatial blocks (6 by 6 grid), with all blocks the same size, and each overlapping neighboring blocks by 10 pixels. Within each block, the motion estimates described above were linearly interpolated to estimate motion within the block, and this offset was applied to correct each frame. After applying this correction offset, there was no apparent residual motion within the block. Within each block, the shapes of active cells were identified using constrained nonnegative matrix factorization [37]. Because adjacent blocks overlapped, some cells were identified more than once. Two identified cells were considered duplicates if their shapes exhibited a Pearson’s correlation exceeding 0.8, and the cell with a smaller spatial extent was removed. Time courses were median-filtered (length 10), and thresholded by zeroing time points below a certain threshold (4 times the robust standard deviation). All subsequent analysis steps were performed using the same procedures as other datasets.

### 4.5. Computing place fields

The spatially-averaged activity was computed by dividing the track into 10 cm spatial bins, averaging the activity that occurred when the mouse was in each bin, then smoothing over space by convolving with a Gaussian kernel (radius 20 cm), with the smoothing kernel wrapping at the edges of the track.

Whether a cell exhibited a spatially-modulated field was defined by how much information its activity provided about linear track position [48]. For each cell, the information I was computed as 
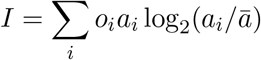

where *o_i_* is the probability of occupancy in spatial bin *i*, *a_i_* is the smoothed mean activity level (Δ*F/F*) while occupying bin *i*, and *ā* is the overall mean activity level. This value was compared to 100 shuffles of the activity (each shuffle was generated by circularly shifting the time course by at least 500 frames, then dividing the time course into 6 chunks and permuting their order). If the observed information value exceeded the 95th percentile of shuffle information values, its field was considered spatially-modulated.

The COM of each spatially-modulated cell was computed by transforming the spatially-averaged activity to polar coordinates, where *θ* was the track position and *r* was the average activity amplitude at that position. The two-dimensional center of mass of these points was computed, and their angle was transformed back to track position to yield the COM location. No special treatment was given to cells that might have multiple fields.

Because the end of the track was continuous with the beginning, its topology was a circle rather than a line segment. To accommodate statistical tests and fits designed for a linear topology, COM locations were re-centered on the region of interest. For Hartigan’s Dip Test, COM locations were centered at 266 cm. For fitting Gaussian distributions to the excess density (see below), COM locations were centered at the reward location.

Reward-associated cells were typically chosen by identifying with COMs located within 25 cm of the reward (before or after). Reward-predictive cells were defined as reward-associated cells with a COM located prior to the reward. Place cells were defined as cells with a spatial field that were not reward-associated. It should be noted that in most cases (e.g. Figures 3-5) a set of putative reward-associated cells selected based on COM location likely contained some place cells that coincidentally exhibited fields near the reward. Though the analysis of Figure 2 showed that reward-associated cells composed a separate class, the identity of given cell active near reward was ambiguous, since it could not be determined whether it came from the distribution of place cells or true reward-associated cells.

### 4.6. Computing activity correlation across environments

To identify how similarly the entire recorded ensemble encoded position on track A and track B during condition AB, the population vector correlation was computed. For each cell, the spatially-averaged and smoothed activity was computed as described above. Since track B was shorter than track A, only the first 250 cm of track A was used. The average activity values of every cell at every position were treated as a single vector, and the Pearson’s correlation was computed between the vector for track A and the vector for track B.

### 4.7. Fitting COM density in condition AB

The density of COMs on each track was fit with a mixture distribution that combined a uniform distribution and a Gaussian distribution, i.e.

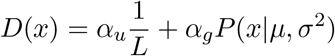

where *D*(*x*) is the total COM density at track location *x*, *α_u_* is the fraction of cells that are uniformly distributed, *L* is the track length, *α_g_* = 1 − *α_u_* is the fraction of cells that compose the excess density near reward, and *P*(·|*μ*, *σ*^2^) is the probability density function for a Gaussian distribution with mean *μ* and variance *σ*^2^. A maximum likelihood fit to the observed COMs was used to estimate the *α* coefficients as well as the parameters of the Gaussian.

The four Gaussian fit parameters (one pair for each track; 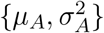 and 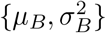) were then used to generate the joint probability densities that would be predicted by hypotheses H1 and H2 for COM locations on the two tracks. Based on these, a mixture distribution was fitted to the observed COMs. Again a maximum likelihood fit was used to estimate the fraction of cells in each component of the mixture distribution:

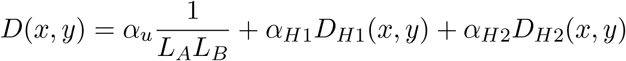

where *D*(*x, y*) was the probability of observing a cell with a COM on track A at location *x* and a COM on track B at location *y*, *α_u_* was the fraction of cells that remapped according to a uniform distribution, *α*_*H*1_ and *α*_*H*2_ were the respective fractions of cells that remapped according to H1 and H2, and the following constraint was applied:

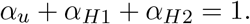

The H1 and H2 distributions were

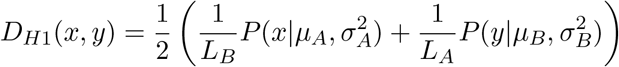

and

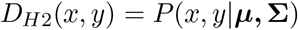

with

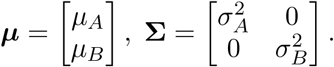

The shape of the H1 and H2 distributions are shown in rough schematic form in Figure 2e. The confidence interval for the parameters of the fit was generated by 1,000 bootstrap resamplings of the observed COM locations.

The fit to the joint density only used cells with a spatial field on both tracks (“dual-track cells”). For cells with a field on only one track (“single-track cells”), remapping could not be followed across environments. However, the identity of those cells (place cells or reward-associated cells) could be inferred by incorporating an additional assumption: that among place cells with a spatial field on at least one track, a random cross-section would also have a field on the other track. This assumption postulated that, for example, all place cells with a field on track A were equally likely to have a field on track B, regardless of their track A field location. This assumption, based on findings of independence across environments [24], implied that single-track and dual-track place cells would exhibit the same COM density. After establishing that among dual-track cells the excess density contained no place cells, it followed that among single-track cells the excess density contained no place cells, and thus on each track the excess density was composed exclusively of reward-associated cells (Supplemental Figure 2).

### 4.8. Identification of slowing, walking bouts, and rest events

Different behavioral states were defined based on movement speed, as detailed below. Graphical illustrations of these states are shown in Supplemental Figure 5.

Instantaneous movement speed was computed as follows. The time course of position was resampled to 30 Hz, and teleports were compensated to compute the total distance travelled. This trace was temporally smoothed using a Gaussian kernel with a radius 2 of samples, then the difference between adjacent time points was computed and smoothed in the same way to yield instantaneous movement speed.

Pre-reward walking bouts began at the moment speed dropped below a mouse-specific threshold for the last time prior to reward delivery. Thresholds were chosen manually by examining typical running speed from sessions late in training. This threshold distinguished “running” from “walking”, with the moment of transition defined as “slowing”. If speed did not fall below threshold at least 5 cm before reward, the mouse was not considered to have slowed prior to reward. This threshold was applied because brief (<5 cm) walking bouts occurred throughout the track at approximately uniform density (not shown), and they might have spuriously overlapped the reward zone even if the mouse was not aware of the current reward location.

Mice sometimes walked slowly at locations that were not immediately prior to the reward, and these unrewarded walking bouts were defined slightly differently. First, candidate bouts were identified as times during which speed was lower than half the threshold. The beginning of the bout was defined as the moment when speed fell below threshold, and they ended when speed rose above half the threshold for the last time prior to rising above the full threshold. If the mouse traversed at least 15 cm during this period, it was considered a walking bout. Walking bouts beginning less than 25 cm after reward were excluded to avoid the periods when mice ramped up their speed prior to running to the next reward. Walking bouts that did or did not span the non-current reward location were categorized separately.

Rest events were defined similarly to walking bouts, with three additional or modified criteria: speed fell to 1 cm/sec or lower at some point during the bout, distance advanced less than 15 cm during the bout, and the bout did not span either reward site.

For comparing the activity during rest events or walking bouts to running, periods of running were defined as the interval between 2 and 5 seconds prior to a walking bout, and only at time points during which speed was at least 20% over threshold.

### 4.9. Percentile Correlation

For each cell, the degree of correlation between activity and speed was quantified with a shuffle test. Values of the spatially-binned speed and activity (as depicted in panels 3f,g) were treated as vectors, and the Pearson’s correlation between them was computed. Equivalent values were also computed for a shuffle distribution, in which activity was randomly assigned to different traversals. If activity tended to occur only after the mouse slowed, the observed correlation would be lower than the shuffle distribution.

Importantly, the percentile correlation was not treated as a p-value. Applying a statistical test of significance to the entire population would require a correction factor for multiple comparisons, a more stringent test that might exclude many cells. Instead, the percentile correlation was treated as a general score, with the null expectation that, for example, 5% of cells would exhibit a value of 5th percentile or less.

The polarity (positive or negative) of the observed correlation was not considered, since it was not necessarily informative about whether the activity of a cell was related to slowing. The correlation could be negative even for a cell that was not related to slowing, and it could be positive for a cell that was precisely aligned to slowing.

### 4.10. Density of Slowing-Correlated Cells

The aim of this measurement was to determine whether slowing-correlated cells occurred more frequently than chance among the populations of place cells and reward-associated cells. Only cells with a spatial field during both contexts in condition A_end_A_mid_ were included, and each cell was assigned to a spatial bin based on it how remapped (50cm bin width, bin edges offset by 16cm to align with reward location). The following analysis was performed separately for cells that were slowing-correlated during A_end_ (Figure 3i) and during A_mid_ (Supplemental Figure 4b).

To estimate the density of slowing-correlated cells in each bin, the numerator was the number of slowing-correlated cells, and the denominator was the number of cells with sufficient activity (a transient onset within 100 cm before reward on at least ten trials). Under the null hypothesis, the density of slowing-correlated cells would be 0.05 in each bin. Because in some bins the numerator and denominator were very small (e.g. 1/3), the maximum likelihood estimate of density (e.g. 0.33) would not reflect the high level of uncertainty. Therefore the estimated density was taken as the lower bound of the 95% confidence interval for a binomially-distributed variable (e.g. 0.0084). The densities of each bin were then smoothed with a Gaussian kernel (radius 0.8 bins), with the convolution wrapping at the edges of the track. Regions in which this spatially-smoothed lower bound estimate exceeded 0.05 were assumed to contain a greater fraction of slowing-correlated cells than predicted by the null hypothesis.

### 4.11. Nonparametric Analysis of Remapping among Slowing-Correlated Cells

This analysis had the same aim as computing the density of slowing-correlate cells, but it was nonparametric in the sense that it did not use the COM location, nor did it explicitly calculate the density. All cells were included that exhibited a spatially-modulated field during the context under consideration (A_end_ or A_mid_). For each cell, activity on each traversal was spatially binned (10 cm bin width), averaged across traversals, and the average was normalized to have a sum of 1. To ensure this reflected the steady-state activity within each block, the first three traversals after block transitions were excluded. The averages of all slowing-correlated cells were combined by taking the mean across cells in each spatial bin. To estimate the baseline fluorescence level for comparison, the same procedure was applied to all cells with a spatial field that were not slowing-correlated.

### 4.12. Time of activity relative to slowing

Activity was binned on each trial (200 msec width) relative to slowing time, averaged across trials, and smoothed by convolving with a Gaussian kernel (s.d. 0.1 sec). The point when this trace assumed its maximum was taken as the time of peak activity for each cell. For Figure 4c, cells were only included if they were active in the 10 seconds prior to slowing on at least 5 traversals of each context.

### 4.13. Combining speed and activity across trials

Image acquisition rates varied across datasets (typically near 12 Hz), and within each dataset the estimate of instantaneous speed (described above) was subsampled to match the frame rate to be sure speed was precisely aligned to activity. To combine across datasets, speed and activity time courses were resampled to 10 Hz (MATLAB “resample” command).

For Figure 5c, a subset of bouts was chosen to ensure the average speed profiles were as similar as possible for bouts before the current and non-current reward. For condition A_end_A_mid_, a bout before the non-current reward was excluded if normalized speed fell below 0.3 in the time interval [−2 −0.5] seconds relative to slowing, or if it rose above 0.8 in [3 4]. For condition AB, a bout before the non-current reward was excluded if normalized speed fell below 0.6 in the time interval [−2 −1]seconds relative to slowing, or if it fell below 0.2 in [0 1] or below 0.01 in [2 3]; and bouts before the current reward were excluded if normalized speed fell below 0.5 in [−2 −1] or rose above 2 in [−2 −1]. These parameters were chosen manually to maximize similarity of speed profiles, thus providing a controlled comparison of activity levels.

### 4.14. Control for selection bias when classifying reward-predictive cells

Reward-predictive cells were defined based on exhibiting an average activity location prior to the current reward, a definition that might bias the population to exclude cells that were active prior to the non-current reward. To overcome this bias, a collection of putative reward cells was selected based on being slowing-correlated (see above), a definition that would not tend to include or exclude cells that were active on other parts of the track. Nevertheless, slowing-correlated cells were biased to include cells that were frequently active near the current reward site. Therefore, for the analysis of Supplemental Figure 7, the activity of these cells was only compared on the other part of the track. For example, if putative reward-predictive cells were selected for being slowing-correlated when mice approached the reward at 366 cm, only their activity preceding 166 cm was used: their activity when approaching the current reward was measured during A_mid_, and their activity when approaching the non-current reward was measured during A_end_. This ensured that the activity being compared was independent of the activity used for cell selection.

## 5. Acknowledgements

This work was supported by NIH NRSA 1F32NS077840-01A1 (JLG) and NIH NIMH R01 2R01MH083686-6 (JLG & DWT). Transgenic mice were made by the Genetically-Encoded Neuronal Indicator and Effector (GENIE) Project at the HHMI’s Janelia Research Campus. We thank D. Aronov, M. Harnett, and A. Charles for comments on the manuscript.

## 6. Author Contributions

JLG and DWT designed the experiments. JLG performed the experiments and analyzed the data. JLG and DWT wrote the paper.

